# Role of 1’-Ribose Cyano Substitution for Remdesivir to Effectively Inhibit Nucleotide Addition and Proofreading in SARS-CoV-2 Viral RNA Replication

**DOI:** 10.1101/2020.04.27.063859

**Authors:** Lu Zhang, Dong Zhang, Xiaowei Wang, Congmin Yuan, Yongfang Li, Xilin Jia, Xin Gao, Hui-Ling Yen, Peter Pak-Hang Cheung, Xuhui Huang

## Abstract

COVID-19 has recently caused a global health crisis and an effective interventional therapy is urgently needed. SARS-CoV-2 RNA-dependent RNA polymerase (RdRp) is a promising but challenging drug target due to its intrinsic proofreading exoribonuclease (ExoN). Remdesivir targeting SARS-CoV-2 RdRp exerts high drug efficacy *in vitro* and *in vivo*. However, its underlying inhibitory mechanisms remain elusive. Here, we performed all-atom molecular dynamics simulations with an accumulated simulation time of 24 microseconds to elucidate the molecular mechanisms underlying the inhibitory effects of Remdesivir. We found that Remdesivir’s 1’-cyano group of possesses the dual role of inhibiting nucleotide addition and proofreading. The presence of its polar 1’-cyano group at an upstream site in RdRp causes instability and hampers RdRp translocation. This leads to a delayed chain termination of RNA extension, which may also subsequently reduce the likelihood for Remdesivir to be cleaved by ExoN acting on the 3’-terminal nucleotide. In addition, our simulations suggest that Remdesivir’s 1’-cyano group can also disrupt the cleavage active site of ExoN via steric interactions, leading to a further reduced cleavage efficiency. Our work provides plausible molecular mechanisms on how Remdesivir inhibits viral RNA replication and may guide rational design for new treatments of COVID-19 targeting viral replication.

## INTRODUCTION

The 2019 novel coronavirus (COVID-19 coronavirus (CoV) or severe acute respiratory syndrome coronavirus 2 (SARS-CoV-2)) has spread rapidly to cause serious outbreaks and eventually a global pandemic. As of Sept 2020, COVID-19 has been reported in 213 countries and almost 26 million cases have been confirmed, resulting in more than 870,000 deaths (1). The new CoV has been declared a global emergency by the World Health Organization, as the outbreak continues to spread globally and no clinically approved interventional therapy is currently available to curb this health crisis.

Antiviral agents are urgently needed to treat COVID-19 patients. Unfortunately, it could take years to develop new interventions, including therapeutic antibodies, cytokines, nucleic acid-based therapies, and vaccines. Hence, repurposing clinically approved or investigative drugs for other diseases provides a promising approach to develop COVID-19 treatments. CoV has a capsid that envelops the single-stranded RNA genome (2). Three structural surface proteins are shown to be associated with the capsid: viral membrane, envelope, and the spike protein (3). Because SARS-CoV-2 and SARS-CoV share 89% sequence identity (4), chemical compounds and monoclonal antibodies that target SARS-CoV surface proteins or interrupt its binding to host viral receptor have been under investigation for treating SARS-CoV-2 (5-7). In addition, new vaccines targeting the viral surface antigens are under intense development for the prevention of COVID-19. Unfortunately, drugs and vaccines with inhibitory mechanisms targeting surface receptors may not be effective. This is due to the constant evolution of surface receptors to acquire drug resistance and evade host immune response (8). In contrast, viral RNA-dependent-RNA polymerase (RdRp) is a protein that is deeply buried inside the viral capsule and is functionally conserved for viral replication, rendering it resistant to the propagation of drug-resistant virus. Thus, RdRp serves as a promising drug target for virus infections (9). Indeed, inhibitors targeting polymerase of SARS-CoV-2, such as Remdesivir and Favipiravir, are currently at phase 3 clinical trials.

Compared with other viral RdRps such as influenza virus and rhinovirus, SARS-CoV-2 RdRp is more challenging for drug development due to its intrinsic proofreading exoribonuclease (ExoN) (10). Non-structural protein (nsp) 12, along with cofactors nsp8 with nsp7, is involved in nascent RNA priming and nucleotide addition (11-19) (Figure 1A and 1C). Nucleotide analogues that inhibit nucleotide addition in nsp12 supposedly should inhibit RNA replication. However, in SARS-CoV-2, ExoN of nsp14 has been shown to play a pivotal role in proofreading (20) and can counteract the efficacy of the nucleotide analogues by excision (10,21-23) (Figure 1B and 1D). For example, the nucleotide analogue Ribavirin is being incorporated in the nascent RNA to inhibit nucleotide addition of SARS-CoV RdRp. However, Ribavirin is readily excised from nascent RNA by nsp14, which may explain its limited efficacy *in vivo* (16).

**Figure 1.**
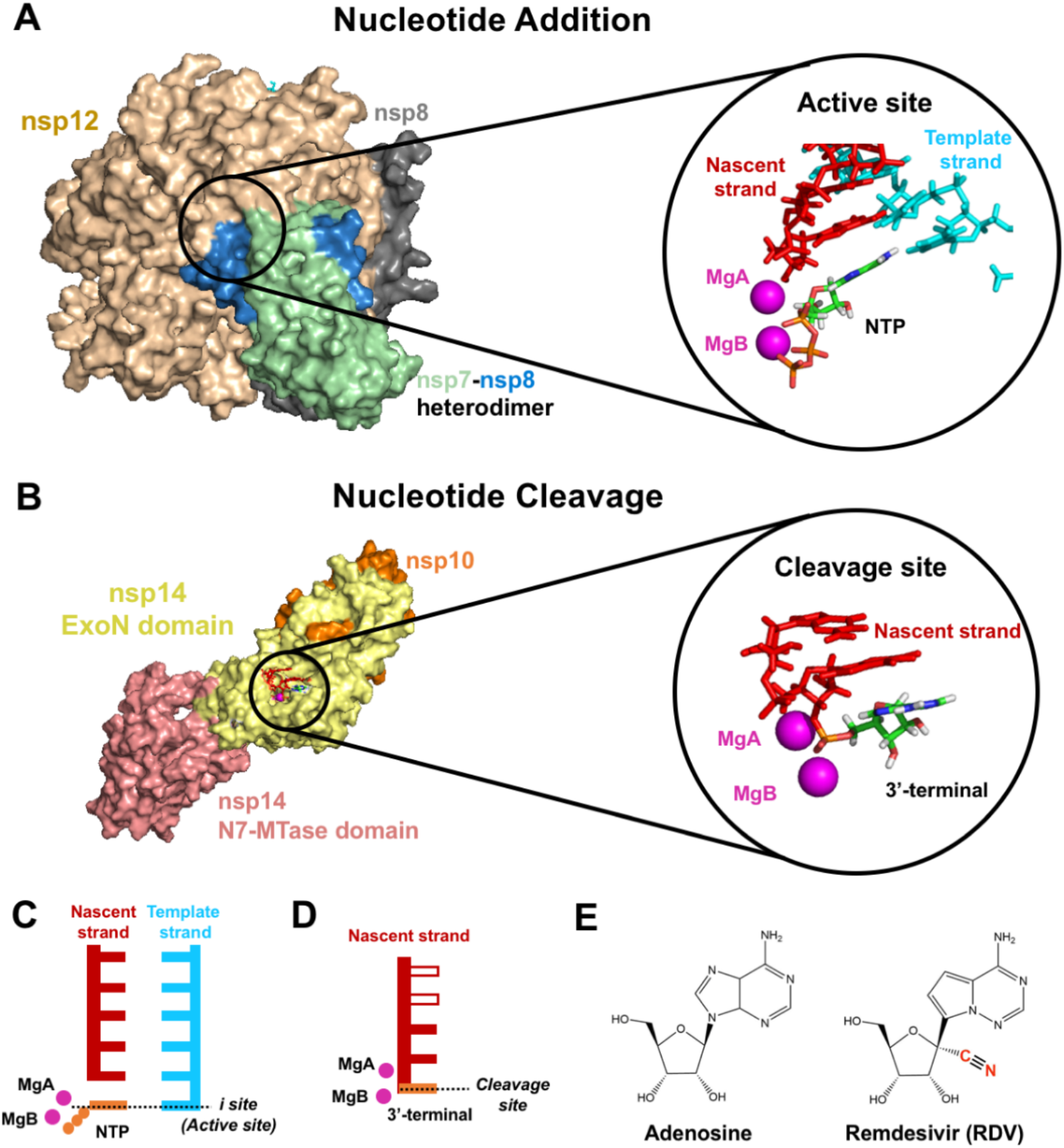
Structural model of SARS-CoV-2 RdRp and ExoN for investigating the inhibitory mechanism of Remdesivir (RDV). (**A**) Model of nsp12-nsp7-nsp8 complex for nucleotide addition. The active site is circled and amplified on the right panel. The nascent and template strands are colored in red and cyan, respectively. NTP (in orange) is bound at the active site and two Mg^2+^ ions are shown in magenta spheres. (**B**) Model of nsp14-nsp10 complex, including ExoN domain for nucleotide cleavage. The cleavage site is circled and amplified on the right. Three nucleotides are modelled, including the 3’-terminal site used for modelling RDV. Magenta spheres represent two Mg^2+^ ions for cleavage. (**C**) Cartoon model of the active site in RdRp. (**D**) Cartoon model of the cleavage site in ExoN. The three terminal nucleotides used in our model are represented by color-filled rectangles. The ones not included in our model are represented by empty rectangles. (**E**) Chemical structures of adenosine and RDV in the nucleoside form. The 1’-cyano group of RDV is highlighted in red.

Remdesivir is a promising drug candidate to treat COVID-19 infection (24-26). It has been shown to be effective as compassionate-use basis to treat patients hospitalised with COVID-19 (26), including those suffering from pneumonia (24). Subsequently, as many as eight clinical trials on using Remdesivir to treat COVID-19 have been conducted (27-29). Remdesivir has shown to be a potent inhibitor of CoV replication in MERS-CoV, SARS-CoV, and circulating human CoVs (22,30). Even though Remdesivir is a nucleotide analogue incorporated by RdRp (Figure 1E), it exerts superior antiviral activity over other nucleotide analogues because the rate of incorporation of Remdesivir to nascent RNA by nsp12 is higher than that of its cleavage by nsp14 ExoN (22). Recently, enormous amount of efforts has been placed in understanding the molecular basis of Remdesivir’s inhibitory mechanisms on RNA synthesis (15,17,18,31-35). Due to the high sequence similarity between SARS and SARS-CoV-2, the nsp12-nsp7-nsp8 and nsp14-nsp10 of SARS-CoV serve as reliable models to study the mechanisms of RNA replication of SARS-CoV-2. Furthermore, the cryo-EM structures of SARS-CoV-2 nsp12-nsp7-nsp8 have been recently solved (15,17-19), with the catalytic domain of nsp12 showing very high structural similarity to that of SARS-CoV (13,16). Biochemical assays have shown in MERS CoV (31) and recently in SARS-CoV-2 (17,19,32) that Remdesivir causes “delayed chain termination” in RNA synthesis, wherein several rounds of nucleotide additions can still proceed after Remdesivir incorporation prior to completed termination. However, the molecular mechanisms underlying this delayed termination induced by Remdesivir remain largely elusive. Furthermore, it is unclear how Remdesivir can evade the proofreading activity of SARS-CoV-2 to prevent itself from being cleaved by ExoN.

To elucidate the mechanisms of Remdesivir inhibition on RdRp and ExoN, we conducted an aggregation of 24 μs all-atom molecular dynamics (MD) simulations of the nsp12-nsp7-nsp8 and nsp14-nsp10 complexes in SARS-CoV-2. For RdRp, our MD simulations show that Remdesivir at the 3’-terminal does not impact the efficiency of nucleotide addition in nsp12. More interestingly, we found that Remdesivir at an upstream site (*i+3* site) can block translocation via interactions formed by its 1’-cyano group, thus resulting in delayed chain termination. As the inhibitory effect is exerted by the Remdesivir at three nucleotides upstream rather than at the 3’-terminal of the nascent RNA, this will also reduce the likelihood for the Remdesivir to be cleaved by ExoN, which only acts on the 3’-terminal nucleotide. Furthermore, we found that the relatively bulky 1’-cyano group on the ribose of Remdesivir introduces steric interactions in ExoN and effectively disrupts the stability of the cleavage active conformation, and thus effectively inhibits the exonuclease activity of ExoN. Our work has underlined the important role of the 1’-cyano substitution in Remdesivir in exerting the inhibitory effect on nucleotide addition and proofreading, providing guidance for the rational design of nucleotide analogues targeting at SARS-CoV-2 RdRp.

## MATERIAL AND METHODS

### Structural model of nsp12-nsp7-nsp8 complex

The cryo-EM structure of nsp12-nsp7-nsp8 complex (PDBID: 6NUR) (13) and the norovirus holo-RdRp (PDBID: 3H5Y (36)) were used as the structural basis to construct the nCoV RdRp containing dsRNA with adenosine triphosphate (ATP) bound at the active site (see SI Section 1.1 for details). The protonation states of histidine residues were predicted using propka3.0 module (37) in the pdbpqr2.2.1 (38) package, followed by manual investigation to ensure that the coordination with the zinc ion can be formed. The whole complex was placed in a dodecahedron box with the box edges at least 12 Å away from the complex surface. The box was filled with TIP3P water molecules (39), and enough counter ions were added to make the whole system neutral.

### Structural model of nsp14-nsp10 complex

The crystal structure of nsp10-nsp14 complex of SARS-CoV (PDBID: 5C8S (40)) serves as the structural basis to construct the nsp10-nsp14 of SARS-CoV-2 (see SI Section 1.2 for details). Missing residues in nsp14 were modelled by modeller9.21 (41). Single-strand RNA containing three nucleotides were modelled by structural alignment to the cleavage site of the proofreading domain of DNA polymerase I Klenow fragment (PDBID: 1KLN (42)) and the ε-subunit of DNA polymerase III (PDBID: 1J53 (43)) because their cleavage sites share similar architectures (40). The protonation states of histidine residues were estimated by propka3.0 module (37) of the pdbpqr2.2.1 (38) package. The complex was solvated with TIP3P water in a dodecahedron box, the edge of which was at least 12 Å away from the complex surface. Sufficient counter ions were added to neutralize the system.

### MD setup and parameters

We used the amber99sb-ildn force field (44) to simulate protein and nucleotides. The force field parameters for Remdesivir in the nucleoside form were derived based on the existing amber99sb-ildn force field parameters or general amber force field (45,46) (see SI Section 2.1 for details). Partial charges of Remdesivir in the nucleoside form were generated by following the Restricted Electrostatic Potential scheme (47,48). For ATP or Remdesivir in the triphosphate form, parameters for the triphosphate tail were taken from those developed by Meagher *et al*. (49). The system was gradually relaxed before the production simulation under NVT ensemble (T=300 K). V-rescale thermostat (50) was applied with the coupling time constant of 0.1 ps. The long-range electrostatic interactions beyond the cut-off at 12 Å were treated with the Particle-Mesh Ewald method (51). Lennard-Jones interactions were smoothly switched off from 10 Å to 12 Å. The neighbor list was updated every 10 steps. An integration time step of 2.0 fs was used and the LINCS algorithm (52) was applied to constrain all the bonds. We saved the snapshots every 20 ps. All simulations were performed with Gromacs 5.0 (40). The equilibrated system containing wildtype-RNA serves as the structural basis for modelling Remdesivir at the corresponding site in nsp12-nsp7-nsp8 or nsp14-nsp10 complexes to investigate their inhibitory effects. The details of MD simulations are in SI Section 2.2 and 2.3.

### Translocation pathway of Remdesivir from *i*+*3* to *i*+*4* site

We used Climber algorithm (53,54) to generate a pathway for the translocation of Remdesivir from *i+3* to *i+4* site (see SI Section 3 for details). The pre-translocation (pre-T) and post-translocation (post-T) conformations were constructed based on the pre-T model with Remdesivir at *i+3* site and post-T model with Remdesivir at *i+4* site, respectively. We performed a 500-step Climber simulation, in which double stranded RNA was gradually driven from the pre-T state to the post-T state by external energy.

## RESULTS

### Remdesivir at the 3’-terminal of nascent RNA strand does not impact nucleotide addition

Our structural model of SARS-CoV-2 RdRp (nsp12-nsp7-nsp8) has been constructed based on the cryo-EM structure of SARS-CoV RdRp (PDBID: 6NUR (13), see Figure 1A). Template and nascent RNA strands have been constructed by the structural alignment of modelled SARS-CoV-2 RdRp to the norovirus RdRp (PDBID: 3H5Y (36), see Material & Methods and SI Section 1.1 for details). Twenty replicas of 100-ns MD simulations were performed for the RdRp in the post-T state, in which the active site (*i* site) is occupied by ATP (Figure 1A and 1C, see SI Section 2.2.1 for details). To validate this model (named as “wildtype-RNA”), we first examined the stability of the active site by assessing the distance between the *α* phosphate (P*α*) of ATP and O3’ atom of the 3’-terminal nucleotide (Figure 2A). Typically, such distance is required to be below 4 Å to allow efficient catalysis to form the phosphodiester bond (e.g. 3.4 Å in the structure of norovirus RdRp (36) and 3.5∼4.0 Å as suggested by previous computational studies (55)). Consistent with this distance requirement, we found that the distribution of the O3’-P*α* distance peaks at ∼3.5 Å (see the gray curve in Figure 2A). Furthermore, the base-to-base distance between ATP and its base-paired template nucleotide was also measured as an estimation of base-pairing stability (56). We found that such base-to-base distance is most populated at ∼7.5 Å, similar to the distance of 7.4∼7.6 Å as observed in structures of other viral polymerases (36,57) (Figure 2D).

**Figure 2.**
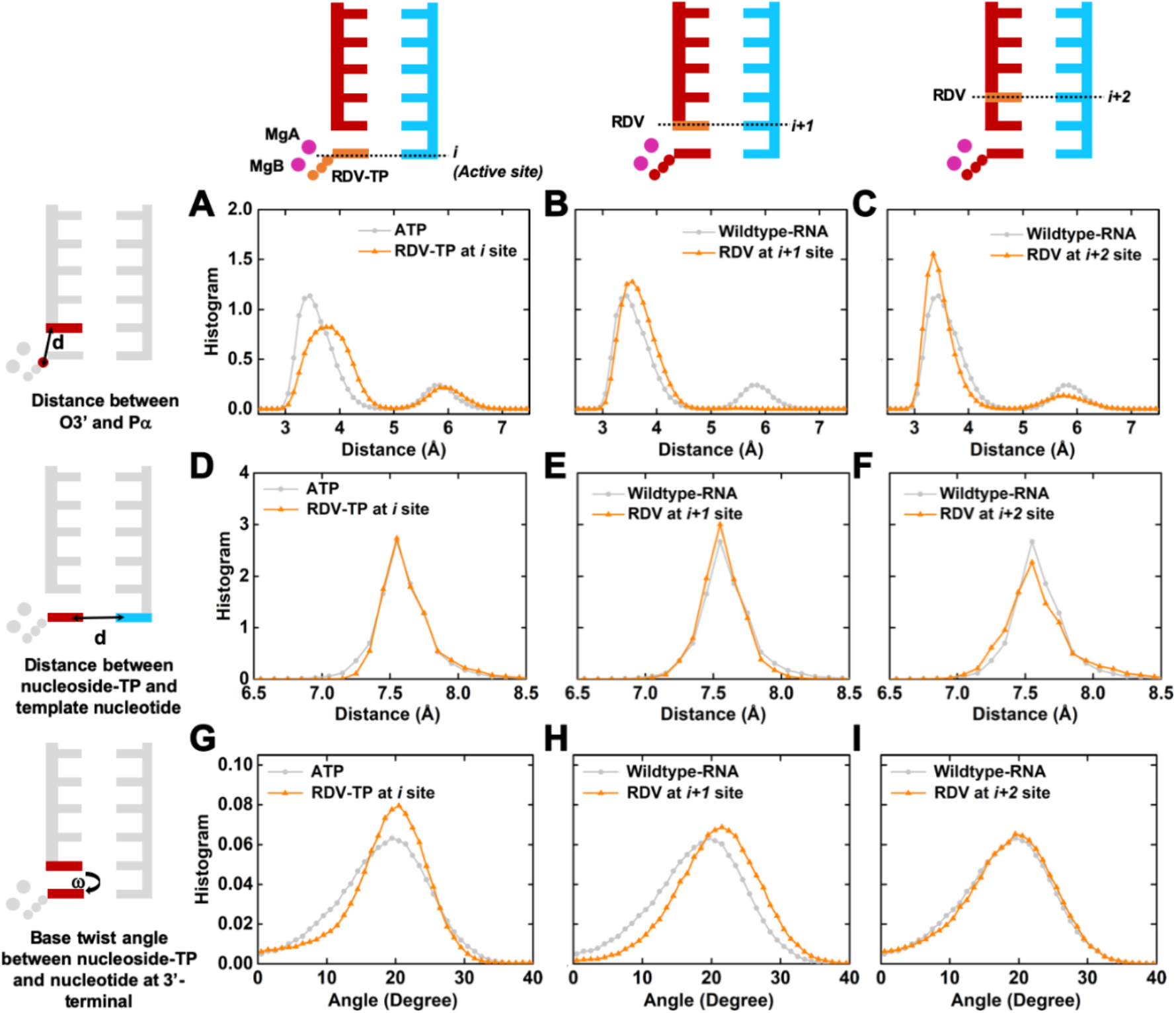
Stability of active site for nucleotide addition in RdRp when RDV is at *i, i+1*, or *i+2* site. The three cartoons in the top panel denote the site where RDV is located (in orange). The three cartoons in the left panel describe the details about the structural features for calculation. For clarification, only the molecules involved in the calculations are colored. Each of the nine histograms is calculated for the structural features (cartoon in the left vertical panel) using the model with RDV at a specific site (cartoon in top horizontal panel). (**A**)-(**C**) Histogram of distance between P*α* of the ATP/RDV-TP and O3’ of the 3’-terminal nucleotide when RDV is at *i* (**A**), *i+1* (**B**), or *i+2* (**C**) site. (**D**)-(**F**) Histogram of distance between the base of ATP/RDV-TP and the base of the corresponding template nucleotide when RDV is at *i* (**D**), *i+1* (**E**) or *i+2* (**F**) site. See SI Section 4.1 for details about the base-to-base distance calculations. (**G**)-(**I**) Histogram of twist angle between the base of ATP/RDV-TP and the base of the 3’-terminal nucleotide when RDV is at *i* (**G**), *i+1* (**H**) or *i+2* (**I**) site. See Supplementary Figure S3 for the illustration of twist angle and SI Section 4.2 for details about the twist angle calculations. For each panel, the histogram for the wildtype-RNA with ATP in the active site is shown in light grey as a reference.

In addition to assessing the active site stability, we further validated our structural model against the two recently solved cryo-EM structures of SARS-CoV-2 RdRp in the post-T state (18,19) (Figure 3A and 3B). The root-mean-square-deviations (RMSDs) of C*α* atoms and nucleotides were used as the metric to examine the structural similarity. We found that the averaged RMSDs are 2.63 Å (PDBID: 6YYT) (18) and 2.66 Å (PDBID: 7BZF) (19) for C*α* atoms, indicating that the overall architecture of our nsp12 model is similar to that of the cryo-EM structures of SARS-CoV-2. The RMSDs of nucleotides between our structural model and the cryo-EM structures were 2.27 Å (PDBID: 6YYT) (18) and 2.57 Å (PDBID: 7BZF) (19), respectively. Finally, we also examined the interactions between upstream RNA and proteins, and found that the interactions observed in the cryo-EM structure are well maintained in our MD simulations (Supplementary Figure S1), further validating our wildtype-RNA model.

**Figure 3.**
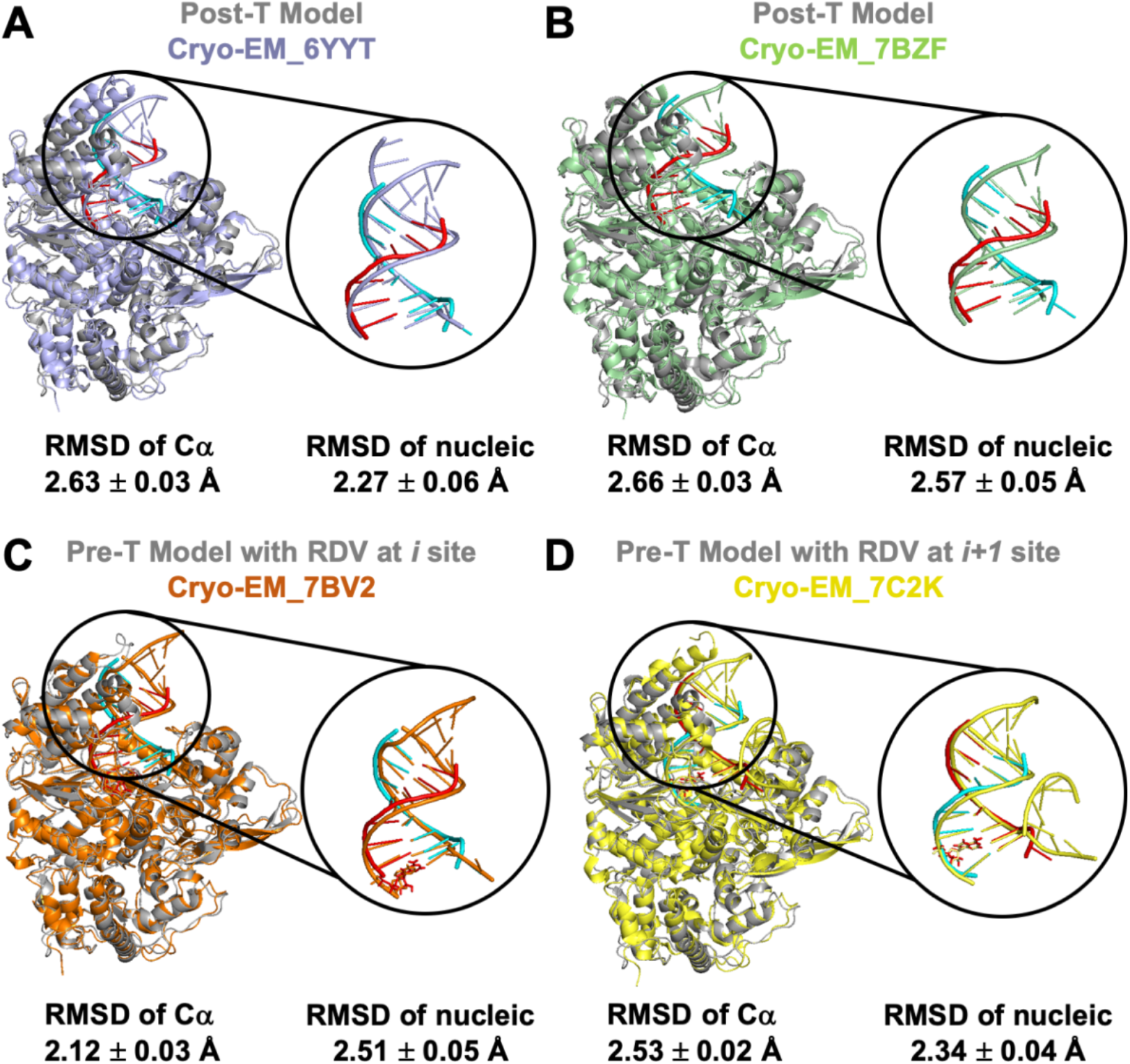
Comparison of our RdRp structural model with the cryo-EM structures. (**A**) Comparison between our model (grey) and cryo-EM structure of RdRp in the post-T state (PDBID: 6YYT, in blue). (**B**) Comparison between our model (grey) and cryo-EM structure of RdRp in the post-T state (PDBID: 7BZF, in green). (**C**) Comparison between our model (grey) and cryo-EM structure of RdRp in the pre-T state with RDV at *i* site (PDBID: 7BV2, in orange). (**D**) Comparison between our model (grey) and cryo-EM structure of RdRp in the pre-T state with RDV at *i+1* site (PDBID: 7C2K, in yellow). In each panel, the template strand (in cyan) and nascent strand (in red) of our model are also compared to those in the cryo-EM structures. RMSD of C*α* and dsRNA are also displayed. See SI Section 4.3 for calculation details.

Next, we examined the if Remdesivir can maintain a catalytically active conformation when it diffuses into the active site. As prodrug Remdesivir (GS-5734) is metabolized into the triphosphate form (Remdesivir-TP) in the cell (Supplementary Figure S2) (58), we performed twenty 100-ns MD simulations with Remdesivir-TP at the active site (*i* site) (see SI Section 2.2.2 for details). Interestingly, we found that the distribution of the critical O3’-P*α* distance in the Remdesivir-TP system is still largely below 4 Å (Figure 2A). Furthermore, the base pairing and base stacking stability (Supplementary Figure S3) are similar to the scenario when ATP is at the active site (Figure 2D and 2G). These results suggest that the Remdesivir-TP can largely maintain a catalytically active conformation, and thus can be added to the 3’-end of the nascent RNA strand. Therefore, Remdesivir satisfies the prerequisite condition to act as a chain-termination inhibitor.

After Remdesivir is added to the RNA chain and is translocated to the adjacent upstream site (*i+1* site), we found that Remdesivir has marginal effect on the efficiency for the next nucleotide addition at the active site. In this case, we also performed twenty 100-ns MD simulations with Remdesivir at 3’-terminal (*i+1* site) to examine if it can inhibit the next ATP incorporation at the active site (see SI Section 2.2.2 for details). We found that the incoming ATP is still stable when Remdesivir is at the *i+1* site. In particular, the P*α*-O3’ distance distribution, as well as the stability of base pairing and base stacking is similar to that observed in the wildtype-RNA (Figure 2B, 2E and 2H). These observations suggest that Remdesivir at *i+1* site is incapable of abolishing the next NTP addition at *i* site.

To examine if Remdesivir at further upstream positions of the nascent RNA strand can interfere with NTP incorporation at the active site, we performed twenty 100-ns MD simulations for each of the systems with Remdesivir at *i+2, i+3* or *i+4* site (see SI Section 2.2.2 for details). The histogram of critical distances for ATP incorporation shows no obvious discrepancy from that of wildtype-RNA (Figure 2C, Supplementary Figure S4A and S4B). Furthermore, the base pairing and base stacking stability are also similar to those in the wildtype-RNA (Figure 2F and 2I, Supplementary Figure S4C-S4F). All these results suggest that Remdesivir at *i+2, i+3* or *i+4* site has negligible impact on nucleotide addition at the active site.

Altogether, we expect that Remdesivir-TP can be incorporated into the nascent strand, and its incorporation does not directly impair NTP addition at the active site. These observations suggest that other mechanisms must exist for Remdesivir to inhibit RdRp function.

### Remdesivir can abolish nucleotide addition in RdRp by hindering translocation

Remdesivir has been proposed as a delayed chain terminator for the RdRps of respiratory syncytial virus (59), Nipah virus (60), and MERS-CoV (31). Recent biochemical experiments for SARS-CoV-2 RdRp also demonstrated delayed chain termination by Remdesivir at physiological concentrations (17,19,32), although nucleotide addition may be terminated earlier when excess amount of Remdesivir is present (17). For MERS-CoV (31) and SARS-CoV-2 (17,19,32) RNA replication, Remdesivir induced delayed chain termination wherein the addition efficiency of the first few nucleotides is maintained. However, nucleotide addition is abruptly abolished at a specific upstream site with a 3-nucleotide delay. The molecular mechanisms underlining this delayed chain termination remain largely elusive.

To elucidate the mechanisms of this delayed termination, we performed MD simulations for RdRp with Remdesivir at different sites (*i* to *i+4* site) of the nascent RNA strand during the translocation process. To obtain statistically meaningful results, we performed twenty 100-ns MD simulations for each of these systems (see SI Section 2.2.2 for details). It is noted that our simulation models of SARS-CoV-2 RdRp in the pre-T state with Remdesivir at *i* or *i+1* site match well with the recent solved cryo-EM structures (Figure 3C and 3D, Supplementary Figure S1) (17,19), providing a validation of our models. To investigate the propensity of translocation, we first inspect how Remdesivir may impact the stability of RdRp complex by examining the probability of hydrogen bonds formed between Remdesivir and the template uracil. As shown in Figure 4A and Supplementary Figure S5, the hydrogen bonds of Remdesivir:U are nearly intact when Remdesivir is at *i, i*+1, or *i*+2 site in both pre-T and post-T states, suggesting that translocation of Remdesivir is allowed in these systems. It is worth mentioning that the interactions between Remdesivir (at *i* site) and surrounding residues in our model are consistent with those in the recently solved SARS-CoV-2 RdRp structure (17) (Supplementary Figure S6). Our observations are also consistent with a recent structural modelling study, suggesting that Remdesivir at the active site (*i* site) cannot hinder translocation (33).

**Figure 4.**
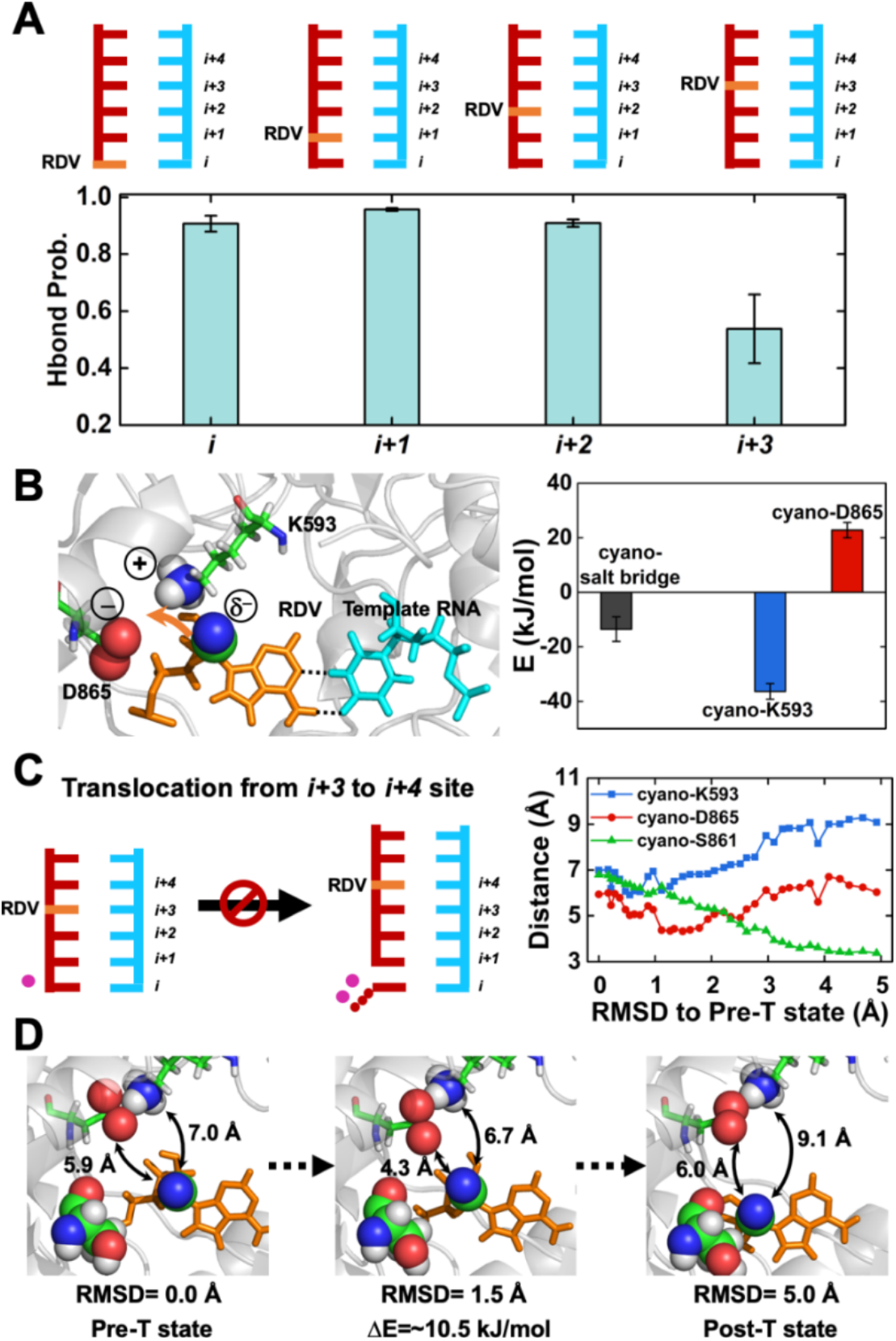
Remdesivir inhibits translocation in RdRp. (**A**) The top panels contain cartoon models of pre-T state with RDV (shown in orange) at *i, i+1, i+2* or *i+3* site. The bottom panel corresponds to the averaged hydrogen bond probability for two hydrogen bonds of the RDV:U base pair when RDV is at *i, i+1, i+2* or *i+3* site (see SI Section 4.4 for calculation details). (**B**) The left panel: A representative MD conformation displaying that the D865^-^-K593^+^ salt bridge attracts the RDV (shown in orange) from its canonical conformation (two hydrogen bonds for RDV:U pair were shown in dashed lines). The 1’-cyano group of RDV, the two oxygen atoms in the carboxyl group of D865 and the quaternary amine group of K593 are shown in spheres. The right panel: Interaction energy between the 1’-cyano group of RDV and D865^-^-K593^+^ salt bridge is shown in black. The interaction energy between 1’-cyano group and D865, as well as that between 1’-cyano group and K593 is shown in red and blue, respectively (see SI Section 4.5 for calculation details). (**C**) Cartoon model showing the translocation with RDV from *i+3* to *i+4* site is inhibited (Left panel). The distance between 1’-cyano group and D865/K593/S861 versus the RMSD of dsRNA to the pre-T state during translocation is shown in the right panel (see SI Section 4.6 and 4.7 for details). (**D**) Representative conformations along the translocation pathway of RDV from *i+3* to *i+4* site.

In sharp contrast, we found that Remdesivir at *i+3* site greatly weakens the hydrogen-bond probability of base pairing with its template uracil (Figure 4A). Further investigations show that the presence of the highly polar 1’-cyano group on Remdesivir will exert strong electrostatic attractions (∼13.6 KJ/mol) with the salt bridge formed by Lys593 and Asp865 of RdRp (Figure 4B). These electrostatic attractions will further pull Remdesivir away from its canonical conformation and cause the destabilization of its base pairing (Figure 4B). Indeed, the hydrogen bonds for the base pairing of Remdesivir:U are subsequently disrupted, rendering the corresponding RdRp complex unstable (Figure 4A). Recently solved cryo-EM structures (17-19) also show that Lys593 and Asp865 are in spatial proximity with 1’-cyano group when Remdesivir was modelled at *i+3* site (Supplementary Figure S7). For translocation of RNA polymerases, a post-T state with the empty active site often has comparable stability with its corresponding pre-T state, and subsequent binding of NTP can greatly stabilize post-T state and favor forward translocation (54). We thus anticipate that translocation of Remdesivir into *i+3* site (post-T) will still be allowed due to the stabilization of NTP binding. However, after NTP incorporation, the unstable complex with Remdesivir at *i+*3 site may hinder the RNA extension. Next, we investigate if this instability induced by Remdesivir at *i+3* site can affect the dynamics of subsequent translocation to *i+4* site by generating a translocation pathway using the Climber algorithm (53,54) (see Material & Methods and SI Section 3 for details). Interestingly, we found that the 1’-cyano group needs to first move closer to the negatively charged Asp865 before it is being translocated away to *i+4* site, while its distance with Lys593 is noticeably larger than that with Asp865 during translocation (Figure 4C). As the electron-withdrawing 1’-cyano moiety exhibits *δ*--anion repulsion with Asp865, the intermediate conformation with reduced distance to Asp865 will experience unfavorable repulsions (∼10.5 kJ/mol raised energy, Figure 4D), and we expect that this may hinder forward translocation (Figure 4C). Altogether, the above observations suggest a delayed termination of three nucleotides as observed in SARS-CoV (32) and MERS-CoV (31).

An alternative explanation was proposed that the steric clash between Ser861 and Remdesivir would inhibit translocation from *i+3* site to *i+4* site (17,32). In our translocation simulation, we found that the 1’-cyano group of Remdesivir constantly approaches Ser861 with their distance monotonically decreased from 7 Å to around 3 Å (Figure 4C). Consistently, the steric clash between Ser861 and the 1’-cyano group of Remdesivir is also observed when we modelled Remdesivir at *i+4* site using the cryo-EM structures (17-19) (Supplementary Figure S8). Due to its spatial proximity, we anticipate that Ser861 also plays an auxiliary role in prohibiting translocation. Based on homology modelling, Shannon *et al*. (33) proposed another mechanism where translocation is inhibited due to the steric clash between Arg858 and Remdesivir’s ribose at *i+4* site. However, our MD simulations with Remdesivir at *i+4* site show a large distance (minimum distance of ∼6.8 Å) between the side chain of Arg858 and 1’-cyano group of Remdesivir. Consistent with our simulations, a large separation (minimum distance of 6.8∼7.9 Å) between Arg858 and the 1’-cyano group of Remdesivir was also observed when we modelled Remdesivir at *i+4* site using the recent cryo-EM structures (17-19) (Supplementary Figure S9). To obtain the translocation pathway, we adopted the Climber method (53,54), in which the RdRp complex is progressively driven from pre-T to post-T state by external energy. To rigorously study the dynamics of large-scale conformational changes during translocation, we need to apply more advanced methods such as Markov State Model (54,61-66), which is beyond the scope of our current study. Finally, a comparison of the sequences between SARS-CoV-2 and other coronaviruses indicates that the Asp865^-^-Lys593^+^ salt bridge is conserved, except that Lys was replaced by another positively charged Arg in two human CoVs (Supplementary Figure S10), implying that our proposed mechanism may be employed by other CoV RdRps.

### Remdesivir can inhibit proofreading

One main challenge for drug design targeting SARS-CoV-2 RdRp is due to the existence of an ExoN domain in nsp14, which has direct interactions with nsp12 (12,40) and performs proofreading function. ExoN can remove the 3’-terminal nucleotide (23) and counteract the efficacy of the inhibitory compounds. Therefore, an additional criterion to be an effective inhibitor for SARS-CoV-2 viral replication is the ability to escape from, or inhibit the cleavage by ExoN.

In SARS-CoV-2, ExoN cannot directly excise the 3’-terminal nucleotide in the RdRp active site, as the channel leading to the active site of SARS-CoV-2 RdRp is too narrow to allow the access of ExoN. Instead, the nascent RNA strand is suggested to backtrack through the channel until it encounters the cleavage site of ExoN (35). Nevertheless, backtracking is energetically disfavored in SARS-CoV-2 RdRp as the RNA base pairs are broken in the upstream, while no base pairs are formed in other regions to compensate the broken base pairs in the backtracking process. A recent structural analysis has demonstrated nsp13 can interfere with the downstream template RNA strand and acts as a motor to trigger the backtracking (35). In addition, any mechanism that can render RdRp to pause or stall can also promote backtracking. In this regard, Remdesivir at *i+3* site can facilitate backtracking by inhibiting forward translocation. Even so, since Remdesivir is not present at the 3’-terminal of the nascent RNA but at three sites upstream (i.e. delayed chain termination), this could also help to reduce the possibility for Remdesivir to be cleaved by ExoN, as ExoN only acts on the 3’-terminal nucleotide. Thus, three nucleotides downstream would need to be excised prior to Remdesivir. In this regard, the delayed chain termination mechanism acted by Remdesivir serves as an effective approach to interfere with its excision by ExoN. However, we also noticed that the efficacy of Remdesivir is sensitive to the existence of ExoN, as a mutant SARS-CoV with deletion of the ExoN gene is more susceptible to Remdesivir inhibition (22). Therefore, besides delayed chain termination mechanism, Remdesivir may adopt a secondary mechanism to further inhibit proofreading. This auxiliary mechanism may be employed when Remdesivir is backtracked and extruded at the cleavage site of ExoN.

To examine the scenario when Remdesivir reaches 3’-terminal of the nascent RNA strand and to evaluate the stability of Remdesivir at the cleavage site of ExoN, we built the structural model for nsp14 bound with its activator nsp10 based on the crystal structure of nsp14-nsp10 of SARS-CoV (PDBID: 5C8S (40)). As previous work suggests that three nucleotide base pairs are melted in the active site of ExoN (42), we modelled a single-stranded RNA containing three nucleotides based on structural alignment to DNA polymerase I Klenow Fragment (42) and DNA Polymerase III ε subunit (43) (see Material & Methods and SI Section 1.2 for details). Twenty 100-ns MD simulations were performed for ExoN with the 3’-terminal of single-stranded RNA occupied by Remdesivir (see SI Section 2.3.2 for details). As a comparison, the same amount of simulations was also performed for the wildtype-RNA, in which adenine nucleotide is occupying the 3’-terminal (see SI Section 2.3.1 for details). To assess the stability of cleavage site, we examined the distance between MgA and the O3’ atom of the second last nucleotide at the 3’-terminal as it is essential for cleavage (67-69) (Figure 5A). This distance is well maintained in majority of the simulations for the wildtype-RNA (Supplementary Figure S11), and the histogram shows the highest peak in the region of 2.0∼2.5 Å (Figure 5A) suitable for cleavage (67-69). By sharp contrast, MD simulations with Remdesivir at 3’-terminal show destabilization of cleavage site in ExoN, with the increased height of the second peak at ∼3.7 Å (Figure 5A and Supplementary Figure S12). Consistently, the base of Remdesivir is bent more significantly than that of adenine nucleotide (Figure 5B), which also suggests that Remdesivir is experiencing unfavorable interactions at the 3’-terminal. Structural analysis indicates that the less stability of Remdesivir in ExoN is caused by its bulky 1’-cyano group. In particular, the 1’-cyano group of Remdesivir exhibits steric clash with the side chain of Asn104, which separates them apart from each other (Figure 5C and Supplementary Figure S13). This observation from MD simulations is consistent with a recent study based on static structural model of nsp14 of SARS-CoV-2, which also proposes that the 1’-cyano group has steric clash with the cleavage site (33).

**Figure 5.**
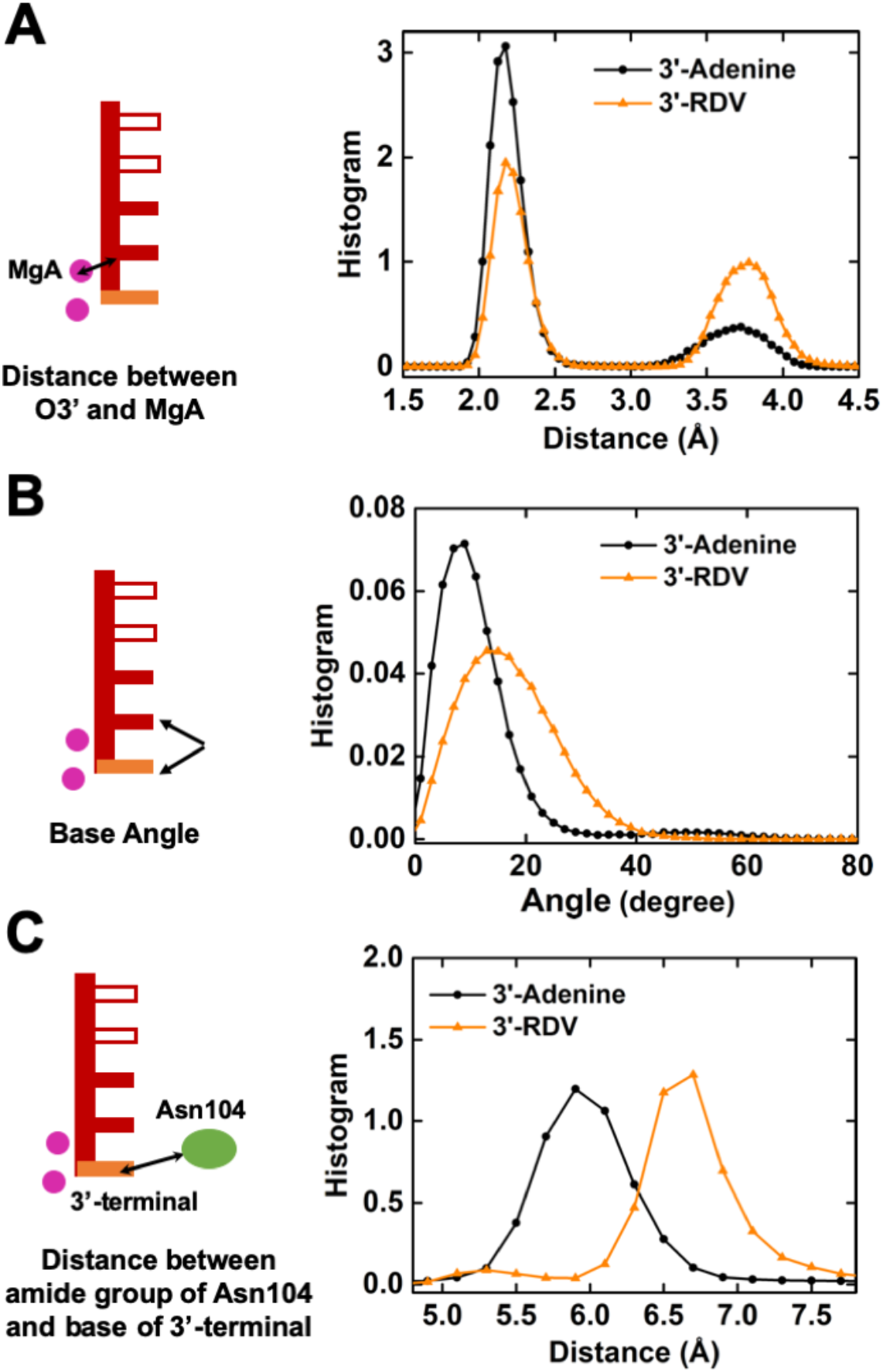
Remdesivir inhibits cleavage in ExoN. (**A**) Histogram of distance between MgA and O3’ atom of the second nucleotide at 3’-terminal. (**B**) Histogram of the base angle between the last two nucleotides at the 3’-terminal. (**C**) Histogram of the distance between the nitrogen atom in the amide group of Asn104 and the base of 3’-terminal (see SI Section 4.8 for details). In (**A**)-(**C**), the histogram for the scenario when RDV is at 3’-terminal is shown in orange, with those when adenine nucleotide is at 3’-terminal shown in black for comparison.

Altogether, the delayed chain termination acted by Remdesivir at *i+3* site of RdRp serves as an effective mechanism to protect Remdesivir from excision by ExoN. The bulky 1’-cyano group introduces extra protection to further reduce the possibility of Remdesivir cleavage in ExoN. The twofold mechanisms collectively enable Remdesivir to be an effective inhibitor for SARS-CoV-2 (70) (Supplementary Figure S14).

## CONCLUSION

In this study, we have elucidated the important role of 1’-cyano group on the ribose in Remdesivir’s inhibition effect targeted RNA replication (Figure 6). Extensive MD simulations were performed for RdRp and ExoN of SARS-CoV-2. Remdesivir can be incorporated into the nascent RNA strand, while its incorporation cannot directly impair the nucleotide addition at the active site. Instead, Remdesivir mainly inhibits nucleotide addition via a delayed chain terminating mechanism, where the growing RNA chain will be terminated once a few nucleotides are added after Remdesivir incorporation. We propose a model where the polar 1’-cyano group of Remdesivir at the upstream *i+3* site will cause instability of the RdRp complex, because the salt bridge (Asp865^-^-Lys593^+^) will pull Remdesivir away from its canonical conformation to break its base pairing. Furthermore, the dynamic transition of translocation from *i+3* site to *i+4* site will be further hindered due to unfavorable anion-*δ*-repulsion between Asp865 and the 1’-cyano group of Remdesivir, as well as the steric effect from Ser861. As a result, Remdesivir leads to a delayed termination of three nucleotides.

**Figure 6.**
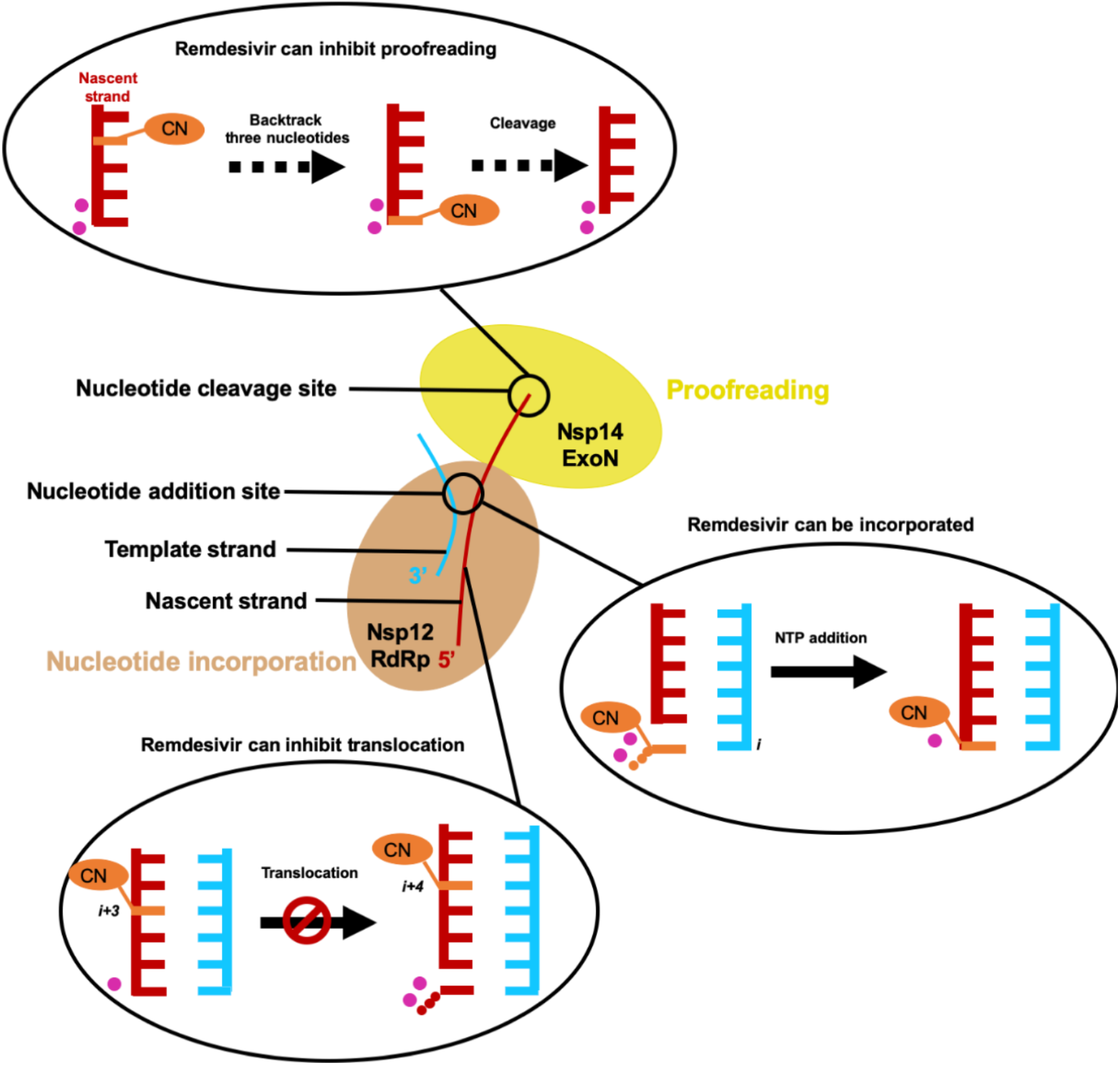
Cartoon model of Remdesivir’s inhibition mechanisms in SARS-CoV-2 viral RNA replication.

ExoN domain of SARS-CoV-2 is responsible for the proofreading function and would cleave the 3’-terminal nucleotide, especially the mismatched one, from the nascent RNA strand (10,21-23). Therefore, the presence of ExoN would reduce the efficacy of nucleotide analogues. Structural modelling indicated that the channel leading to the active site of RdRp is too narrow to allow the ExoN direct access to the 3’-terminal nucleotide in the active site (35). Therefore, Remdesivir at *i+3* site, more upstream than the 3’-terminal, is not possible to be directly cleaved by ExoN. Instead, the nascent RNA strand needs to backtrack three nucleotides until the Remdesivir reaches the 3’-terminal for cleavage (Figure 6). This indicates that the delayed chain termination mechanism can help to protect Remdesivir from cleavage by ExoN. However, previous work by Agostini *et al*. demonstrated that Remdesivir’s inhibitory effect is strengthened on SARS-CoV lacking ExoN (22), indicating Remdesivir can still backtrack to the cleavage site of ExoN. In this scenario, Remdesivir was found to be capable of inhibiting the proofreading function of ExoN. The 1’-cyano group of Remdesivir exhibits steric clash with a nearby residue Asn104, which destabilizes Remdesivir at the cleavage site. Therefore, the delayed chain termination mechanism acted by Remdesivir at *i+3* site of SARS-CoV-2 RdRp plays an essential role in preventing Remdesivir from cleavage by ExoN, and the steric effect that Remdesivir experiences in the cleavage site of SARS-CoV-2 ExoN renders it more resistant to ExoN cleavage, thus facilitating Remdesivir to inhibit RNA synthesis.

Our work provides mechanistic insights at atomistic details on how Remdesivir can effectively inhibit SARS-CoV-2 RNA synthesis. We also revealed that the chemical modification at the ribose of Remdesivir is responsible for the dual inhibitory function to terminate both nucleotide addition and proofreading. This knowledge is valuable for offering guiding principles to maintain the property of Remdesivir to inhibit viral replication, while optimizing Remdesivir’s other drug properties such as preventing its breakdown in the liver (71) and reducing its cytotoxicity (26). Our work also provides a promising general strategy for designing prospective nucleotide analogue inhibitors to treat COVID-19.

## DATA AVAILABILITY

The MD simulation setup files and the MD trajectories are available at https://osf.io/8yhar/?view_only=1983c4aca39e4410bd3f8609f01619c9.

## ACKNOWLEDGEMENT

This research made use of the computing resources of the Supercomputing Laboratory at King Abdullah University of Science and Technology. The National Natural Science Foundation of China Grants [grant number 21733007 and 21803071 to L. Z.]; Shenzhen Science and Technology Innovation Committee [grant number JCYJ20170413173837121 to X. H.]; Hong Kong Research Grant Council [grant number 16307718, 16318816, AoE/P-705/16, and T13-605/18-W to X. H; 16301319 to P.P.C.]; Innovation and Technology Commission [grant number ITCPD/17-9 to X. H.] and the King Abdullah University of Science and Technology (KAUST) Office of Sponsored Research (OSR) [Award No. FCC/1/1976-18, FCC/1/1976-23, FCC/1/1976-26, URF/1/4098-01-01, and REI/1/0018-01-01 to X. G.].

## REFERENCES

1. WHO, Coronavirus disease 2019 (COVID-19): Situation report (2020)

2. Lai, M.M. and Cavanagh, D. (1997), Advances in virus research. Elsevier, Vol. 48, pp. 1–100.

3. Li, F. (2016) Structure, function, and evolution of coronavirus Spike proteins. Annu. Rev. Virol., 3, 237–261.

4. Yuen, K.-S., Ye, Z.W., Fung, S.-Y., Chan, C.-P. and Jin, D.-Y. (2020) SARS-CoV-2 and COVID-19: The most important research questions. Cell Biosci., 10, 40.

5. Vincent, M.J., Bergeron, E., Benjannet, S., Erickson, B.R., Rollin, P.E., Ksiazek, T.G., Seidah, N.G. and Nichol, S.T. (2005) Chloroquine is a potent inhibitor of SARS coronavirus infection and spread. Virol. J., 2, 69.

6. Pascal, K.E., Coleman, C.M., Mujica, A.O., Kamat, V., Badithe, A., Fairhurst, J., Hunt, C., Strein, J., Berrebi, A., Sisk, J.M. et al. (2015) Pre- and postexposure efficacy of fully human antibodies against Spike protein in a novel humanized mouse model of MERS-CoV infection. Proc. Natl. Acad. Sci. U. S. A., 112, 8738–8743.

7. Keefe, B.R., Giomarelli, B., Barnard, D.L., Shenoy, S.R., Chan, P.K.S., McMahon, J.B., Palmer, K.E., Barnett, B.W., Meyerholz, D.K., Wohlford-Lenane, C.L. et al. (2010) Broad-spectrum *in vitro* activity and *in vivo* efficacy of the antiviral protein Griffithsin against emerging viruses of the family coronaviridae. J. Virol., 84, 2511–2521.

8. Kennedy, D.A. and Read, A.F. (2018) Why the evolution of vaccine resistance is less of a concern than the evolution of drug resistance. Proc. Natl. Acad. Sci. U. S. A., 115, 12878–12886.

9. Lythgoe, M.P. and Middleton, P. (2020) Ongoing clinical trials for the management of the COVID-19 pandemic. Trends Pharmacol. Sci., 41, 363–382.

10. Denison, M.R., Graham, R.L., Donaldson, E.F., Eckerle, L.D. and Baric, R.S. (2011) Coronaviruses an RNA proofreading machine regulates replication fidelity and diversity. RNA Biol., 8, 270–279.

11. Imbert, I., Guillemot, J.-C., Bourhis, J.-M., Bussetta, C., Coutard, B., Egloff, M.-P., Ferron, F., Gorbalenya, A.E. and Canard, B. (2006) A second, non-canonical RNA-dependent RNA polymerase in SARS coronavirus. EMBO J., 25, 4933–4942.

12. Subissi, L., Posthuma, C.C., Collet, A., Zevenhoven-Dobbe, J.C., Gorbalenya, A.E., Decroly, E., Snijder, E.J., Canard, B. and Imbert, I. (2014) One severe acute respiratory syndrome coronavirus protein complex integrates processive RNA polymerase and exonuclease activities. Proc. Natl. Acad. Sci. U. S. A., 111, E3900–E3909.

13. Kirchdoerfer, R.N. and Ward, A.B. (2019) Structure of the SARS-CoV nsp12 polymerase bound to nsp7 and nsp8 co-factors. Nat. Commun., 10, 2342.

14. Zhai, Y., Sun, F., Li, X., Pang, H., Xu, X., Bartlam, M. and Rao, Z. (2005) Insights into SARS-CoV transcription and replication from the structure of the nsp7–nsp8 hexadecamer. Nat. Struct. Mol. Biol., 12, 980–986.

15. Gao, Y., Yan, L., Huang, Y., Liu, F., Zhao, Y., Cao, L., Wang, T., Sun, Q., Ming, Z., Zhang, L. et al. (2020) Structure of the RNA-dependent RNA polymerase from COVID-19 virus. Science, 368, 779–782.

16. Ferron, F., Subissi, L., Silveira De Morais, A.T., Le, N.T.T., Sevajol, M., Gluais, L., Decroly, E., Vonrhein, C., Bricogne, G., Canard, B. et al. (2018) Structural and molecular basis of mismatch correction and ribavirin excision from coronavirus RNA. Proc. Natl. Acad. Sci. U. S. A., 115, E162–E171.

17. Yin, W., Mao, C., Luan, X., Shen, D.-D., Shen, Q., Su, H., Wang, X., Zhou, F., Zhao, W., Gao, M. et al. (2020) Structural basis for inhibition of the RNA-dependent RNA polymerase from SARS-CoV-2 by Remdesivir. Science, 368, 1499–1504.

18. Hillen, H.S., Kokic, G., Farnung, L., Dienemann, C., Tegunov, D. and Cramer, P. (2020) Structure of replicating SARS-CoV-2 polymerase. Nature, 584, 154–156.

19. Wang, Q., Wu, J., Wang, H., Gao, Y., Liu, Q., Mu, A., Ji, W., Yan, L., Zhu, Y., Zhu, C. et al. (2020) Structural basis for RNA replication by the SARS-CoV-2 polymerase. Cell, 182, 417–428.

20. Eckerle, L.D., Becker, M.M., Halpin, R.A., Li, K., Venter, E., Lu, X., Scherbakova, S., Graham, R.L., Baric, R.S., Stockwell, T.B. et al. (2010) Infidelity of SARS-CoV nsp14-exonuclease mutant virus replication is revealed by complete genome sequencing. PLoS Pathog., 6, e1000896.

21. Smith, E.C., Blanc, H., Vignuzzi, M. and Denison, M.R. (2013) Coronaviruses lacking exoribonuclease activity are susceptible to lethal mutagenesis: evidence for proofreading and potential therapeutics. PLoS Pathog., 9, e1003565.

22. Agostini, M.L., Andres, E.L., Sims, A.C., Graham, R.L., Sheahan, T.P., Lu, X., Smith, E.C., Case, J.B., Feng, J.Y., Jordan, R. et al. (2018) Coronavirus susceptibility to the antiviral Remdesivir (GS-5734) Is mediated by the viral polymerase and the proofreading exoribonuclease. mBio, 9, e00221–00218.

23. Bouvet, M., Imbert, I., Subissi, L., Gluais, L., Canard, B. and Decroly, E. (2012) RNA 3 ‘- end mismatch excision by the Severe Acute Respiratory Syndrome Coronavirus nonstructural protein nsp10/nsp14 Exoribonuclease complex. Proc. Natl. Acad. Sci. U. S. A., 109, 9372–9377.

24. Holshue, M.L., DeBolt, C., Lindquist, S., Lofy, K.H., Wiesman, J., Bruce, H., Spitters, C., Ericson, K., Wilkerson, S., Tural, A. et al. (2020) First case of 2019 Novel Coronavirus in the United States. N. Engl. J. Med., 382, 929–936.

25. Wang, M., Cao, R., Zhang, L., Yang, X., Liu, J., Xu, M., Shi, Z., Hu, Z., Zhong, W. and Xiao, G. (2020) Remdesivir and Chloroquine effectively inhibit the recently emerged Novel Coronavirus (2019-nCoV) *in vitro*. Cell Res., 30, 269–271.

26. Grein, J., Ohmagari, N., Shin, D., Diaz, G., Asperges, E., Castagna, A., Feldt, T., Green, G., Green, M.L., Lescure, F.-X. et al. (2020) Compassionate use of Remdesivir for patients with severe COVID-19. N. Engl. J. Med., 382, 2327–2336.

27. Goldman, J.D., Lye, D.C.B., Hui, D.S., Marks, K.M., Bruno, R., Montejano, R., Spinner, C.D., Galli, M., Ahn, M.-Y., Nahass, R.G. et al. (2020) Remdesivir for 5 or 10 days in patients with severe COVID-19. N. Engl. J. Med.

28. Beigel, J.H., Tomashek, K.M., Dodd, L.E., Mehta, A.K., Zingman, B.S., Kalil, A.C., Hohmann, E., Chu, H.Y., Luetkemeyer, A., Kline, S. et al. (2020) Remdesivir for the treatment of COVID-19 — Preliminary report. N. Engl. J. Med.

29. Wang, Y., Zhang, D., Du, G., Du, R., Zhao, J., Jin, Y., Fu, S., Gao, L., Cheng, Z., Lu, Q. et al. (2020) Remdesivir in adults with severe COVID-19: a randomised, double-blind, placebo-controlled, multicentre trial. The Lancet, 395, 1569–1578.

30. Sheahan, T.P., Sims, A.C., Graham, R.L., Menachery, V.D., Gralinski, L.E., Case, J.B., Leist, S.R., Pyrc, K., Feng, J.Y., Trantcheva, I. et al. (2017) Broad-spectrum antiviral GS-5734 inhibits both epidemic and zoonotic coronaviruses. Sci. Transl. Med., 9, eaal3653.

31. Gordon, C.J., Tchesnokov, E.P., Feng, J.Y., Porter, D.P. and Götte, M. (2020) The antiviral compound remdesivir potently inhibits RNA-dependent RNA polymerase from Middle East respiratory syndrome coronavirus. J. Biol. Chem., 295, 4773–4779.

32. Gordon, C.J., Tchesnokov, E.P., Woolner, E., Perry, J.K., Feng, J.Y., Porter, D.P. and Gotte, M. (2020) Remdesivir is a direct-acting antiviral that inhibits RNA-dependent RNA polymerase from Severe Acute Respiratory Syndrome Coronavirus 2 with high potency. J. Biol. Chem., 295, 6785–6797.

33. Shannon, A., Le, N.T.-T., Selisko, B., Eydoux, C., Alvarez, K., Guillemot, J.-C., Decroly, E., Peersen, O., Ferron, F. and Canard, B. (2020) Remdesivir and SARS-CoV-2: Structural requirements at both nsp12 RdRp and nsp14 Exonuclease active-sites. Antiviral Res., 178, 104793.

34. Zhang, L. and Zhou, R. (2020) Structural basis of the potential binding nechanism of Remdesivir to SARS-CoV-2 RNA-dependent RNA polymerase. J. Phys. Chem. B, 124, 6955–6962.

35. Chen, J., Malone, B., Llewellyn, E., Grasso, M., Shelton, P.M.M., Olinares, P.D.B., Maruthi, K., Eng, E.T., Vatandaslar, H., Chait, B.T. et al. (2020) Structural basis for helicase-polymerase coupling in the SARS-CoV-2 replication-transcription complex. Cell.

36. Zamyatkin, D.F., Parra, F., Machín, Á., Grochulski, P. and Ng, K.K.S. (2009) Binding of 2′-amino-2′-deoxycytidine-5′-triphosphate to Norovirus polymerase induces rearrangement of the active site. J. Mol. Biol., 390, 10–16.

37. Olsson, M.H.M., Søndergaard, C.R., Rostkowski, M. and Jensen, J.H. (2011) PROPKA3: Consistent treatment of internal and surface residues in empirical pKa predictions. J. Chem. Theory Comput., 7, 525–537.

38. Dolinsky, T.J., Nielsen, J.E., McCammon, J.A. and Baker, N.A. (2004) PDB2PQR: an automated pipeline for the setup of Poisson–Boltzmann electrostatics calculations. Nucleic Acids Res., 32, W665–W667.

39. Jorgensen, W.L., Chandrasekhar, J., Madura, J.D., Impey, R.W. and Klein, M.L. (1983) Comparison of simple potential functions for simulating liquid water. J. Chem. Phys., 79, 926–935.

40. Abraham, M.J., Murtola, T., Schulz, R., Páll, S., Smith, J.C., Hess, B. and Lindahl, E. (2015) GROMACS: High performance molecular simulations through multi-level parallelism from laptops to supercomputers. SoftwareX, 1, 19–25.

41. Webb, B. and Sali, A. (2016) Comparative protein structure modeling using MODELLER. Curr. Protoc. Bioinf., 15, 5–6.

42. Beese, L.S., Derbyshire, V. and Steitz, T.A. (1993) Structure of DNA polymerase I Klenow fragment bound to duplex DNA. Science, 260, 352–355.

43. Hamdan, S., Carr, P.D., Brown, S.E., Ollis, D.L. and Dixon, N.E. (2002) Structural basis for proofreading during replication of the *Escherichia coli* chromosome. Structure, 10, 535–546.

44. Lindorff-Larsen, K., Piana, S., Palmo, K., Maragakis, P., Klepeis, J.L., Dror, R.O. and Shaw, D.E. (2010) Improved side-chain torsion potentials for the Amber ff99SB protein force field. Proteins: Struct., Funct., Bioinf., 78, 1950–1958.

45. Wang, J.M., Wolf, R.M., Caldwell, J.W., Kollman, P.A. and Case, D.A. (2004) Development and testing of a general amber force field. J. Comput. Chem., 25, 1157–1174.

46. Wang, J., Wang, W., Kollman, P.A. and Case, D.A. (2006) Automatic atom type and bond type perception in molecular mechanical calculations. J. Mol. Graphics Modell., 25, 247–260.

47. Woods, R.J. and Chappelle, R. (2000) Restrained electrostatic potential atomic partial charges for condensed-phase simulations of carbohydrates. J. Mol. Struct.: THEOCHEM, 527, 149–156.

48. Wang, J., Cieplak, P. and Kollman, P.A. (2000) How well does a restrained electrostatic potential (RESP) model perform in calculating conformational energies of organic and biological molecules? J. Comput. Chem., 21, 1049–1074.

49. Meagher, K.L., Redman, L.T. and Carlson, H.A. (2003) Development of polyphosphate parameters for use with the AMBER force field. J. Comput. Chem., 24, 1016–1025.

50. Bussi, G., Donadio, D. and Parrinello, M. (2007) Canonical sampling through velocity rescaling. J. Chem. Phys., 126, 014101.

51. Essmann, U., Perera, L., Berkowitz, M.L., Darden, T., Lee, H. and Pedersen, L.G. (1995) A smooth particle mesh Ewald method. J. Chem. Phys., 103, 8577–8593.

52. Hess, B., Bekker, H., Berendsen, H.J.C. and Fraaije, J.G.E.M. (1997) LINCS: A linear constraint solver for molecular simulations. J. Comput. Chem., 18, 1463–1472.

53. Weiss, D.R. and Levitt, M. (2009) Can morphing methods predict intermediate structures? J. Mol. Biol., 385, 665–674.

54. Silva, D.A., Weiss, D.R., Avila, F.P., Da, L.T., Levitt, M., Wang, D. and Huang, X.H. (2014) Millisecond dynamics of RNA polymerase II translocation at atomic resolution. Proc. Natl. Acad. Sci. U. S. A., 111, 7665–7670.

55. Sgrignani, J. and Magistrato, A. (2012) The structural role of Mg2+ Ions in a class I RNA polymerase ribozyme: A Molecular simulation study. J. Phys. Chem. B, 116, 2259–2268.

56. Huang, X., Wang, D., Weiss, D.R., Bushnell, D.A., Kornberg, R.D. and Levitt, M. (2010) RNA polymerase II trigger loop residues stabilize and position the incoming nucleotide triphosphate in transcription. Proc. Natl. Acad. Sci. U. S. A., 107, 15745–15750.

57. Gong, P. and Peersen, O.B. (2010) Structural basis for active site closure by the poliovirus RNA-dependent RNA polymerase. Proc. Natl. Acad. Sci. U. S. A., 107, 22505–22510.

58. Eastman, R.T., Roth, J.S., Brimacombe, K.R., Simeonov, A., Shen, M., Patnaik, S. and Hall, M.D. (2020) Remdesivir: A review of its discovery and development leading to emergency use authorization for treatment of COVID-19. ACS Cent. Sci., 6, 672–683.

59. Tchesnokov, E.P., Feng, J.Y., Porter, D.P. and Götte, M. (2019) Mechanism of inhibition of Ebola virus RNA-dependent RNA polymerase by Remdesivir. Viruses, 11, 326.

60. Jordan, P.C., Liu, C., Raynaud, P., Lo, M.K., Spiropoulou, C.F., Symons, J.A., Beigelman, L. and Deval, J. (2018) Initiation, extension, and termination of RNA synthesis by a paramyxovirus polymerase. PLoS Pathog., 14, e1006889.

61. Klippenstein, S.J., Pande, V.S. and Truhlar, D.G. (2014) Chemical kinetics and mechanisms of complex systems: A perspective on recent theoretical advances. J. Am. Chem. Soc., 136, 528–546.

62. Chodera, J.D. and Noe, F. (2014) Markov state models of biomolecular conformational dynamics. Curr. Opin. Struct. Biol., 25, 135–144.

63. Bowman, G.R., Huang, X.H. and Pande, V.S. (2009) Using generalized ensemble simulations and Markov State Models to identify conformational states. Methods, 49, 197–201.

64. Zhang, L., Pardo-Avila, F., Unarta, I.C., Cheung, P.P.H., Wang, G., Wang, D. and Huang, X.H. (2016) Elucidation of the dynamics of transcription elongation by RNA polymerase II using kinetic network models. Acc. Chem. Res., 49, 687–694.

65. Wang, W., Cao, S.Q., Zhu, L.Z. and Huang, X.H. (2018) Constructing Markov State Models to elucidate the functional conformational changes of complex biomolecules. Wiley Interdiscip. Rev.: Comput. Mol. Sci., 8, e1343.

66. Da, L.-T., Pardo-Avila, F., Xu, L., Silva, D.-A., Zhang, L., Gao, X., Wang, D. and Huang, X. (2016) Bridge helix bending promotes RNA polymerase II backtracking through a critical and conserved threonine residue. Nat. Commun., 7, 11244.

67. Lan, P., Tan, M., Zhang, Y., Niu, S., Chen, J., Shi, S., Qiu, S., Wang, X., Peng, X., Cai, G. et al. (2018) Structural insight into precursor tRNA processing by yeast ribonuclease P. Science, 362, eaat6678.

68. Lei, J., Sheng, G., Cheung, P.P.-H., Wang, S., Li, Y., Gao, X., Zhang, Y., Wang, Y. and Huang, X. (2019) Two symmetric arginine residues play distinct roles in *Thermus thermophilus* Argonaute DNA guide strand-mediated DNA target cleavage. Proc. Natl. Acad. Sci. U. S. A., 116, 845–853.

69. Tse, C.K.M., Xu, J., Xu, L., Sheong, F.K., Wang, S., Chow, H.Y., Gao, X., Li, X., Cheung, P.P.-H., Wang, D. et al. (2019) Intrinsic cleavage of RNA polymerase II adopts a nucleobase-independent mechanism assisted by transcript phosphate. Nat. Catal., 2, 228–235.

70. Choy, K.-T., Wong, A.Y.-L., Kaewpreedee, P., Sia, S.-F., Chen, D., Hui, K.P.Y., Chu, D.K.W., Chan, M.C.W., Cheung, P.P.-H. and Huang, X. (2020) Remdesivir, lopinavir, emetine, and homoharringtonine inhibit SARS-CoV-2 replication *in vitro*. Antiviral Res., 104786.

71. Yang, K. (2020) What do we know about Remdesivir drug interactions? Clin. Transl. Sci.

