## Supplemental materials for "Role of 1’-Ribose Cyano Substitution for Remdesivir to Effectively Inhibit Nucleotide Addition and Proofreading in SARS-CoV-2 Viral RNA Replication"

*Supplementary Figures:*

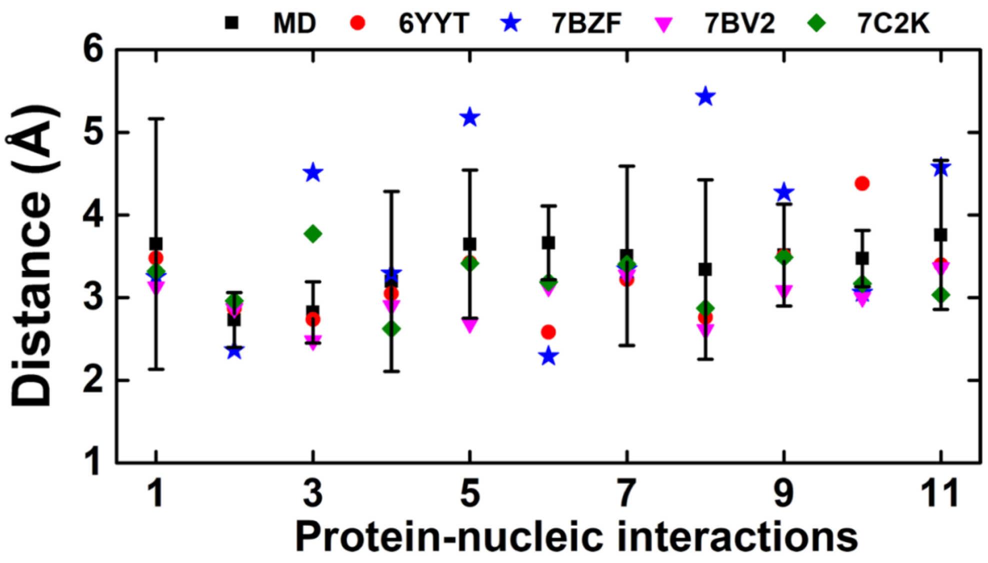


| Index | Chain | Resid | Residue | Interacting atom | Chain | Site | Interacting atom |
|---|---|---|---|---|---|---|---|
| 1 | A | 577 | Lys | NZ | T | *i+3* | phosphate backbone |
| 2 | A | 595 | Tyr | OH | T | *i+6* | phosphate backbone |
| 3 | A | 814 | Ser | OG | P | *i+1* | phosphate backbone |
| 4 | A | 836 | Arg | NH1/NH2 | P | *i+2* | phosphate backbone |
| 5 | A | 858 | Arg | NH1/NH2 | P | *i+4* | phosphate backbone |
| 6 | A | 684 | Asp | O | T | *i+1* | O2' |
| 7 | A | 689 | Tyr | OH | T | *i+2* | O2' |
| 8 | A | 592 | Ser | OG | T | *i+4* | O2' |
| 9 | A | 592 | Ser | OG | T | *i+4* | O3' |
| 10 | A | 759 | Ser | OG | P | *i+1* | O2' |
| 11 | A | 865 | Asp | OD1/OD2 | P | *i+4* | O2' |

**Figure S1.** Validation of our model by comparing protein-nucleotide interactions with cryo-EM structures of SARS-CoV-2 RdRp. The detailed information about the atom pairs used for the calculations is tabulated in the bottom panel. The means and standard deviations of the distances from MD simulations were calculated using all the MD conformations (after removing the first 10ns from each MD trajectory) of RdRp in the post-T state with ATP at *i* site.

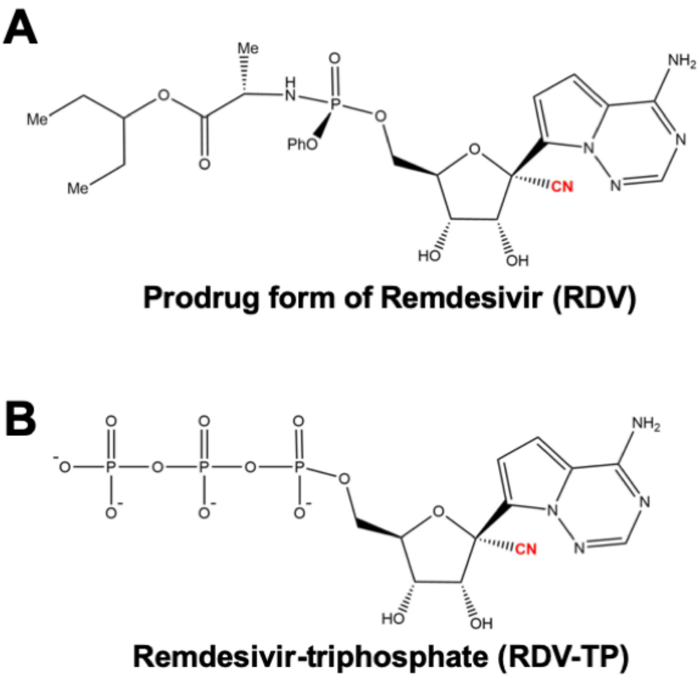


**Figure S2.** Chemical structure of Remdesivir (RDV) in the prodrug form (**A**) and active form (**B**).

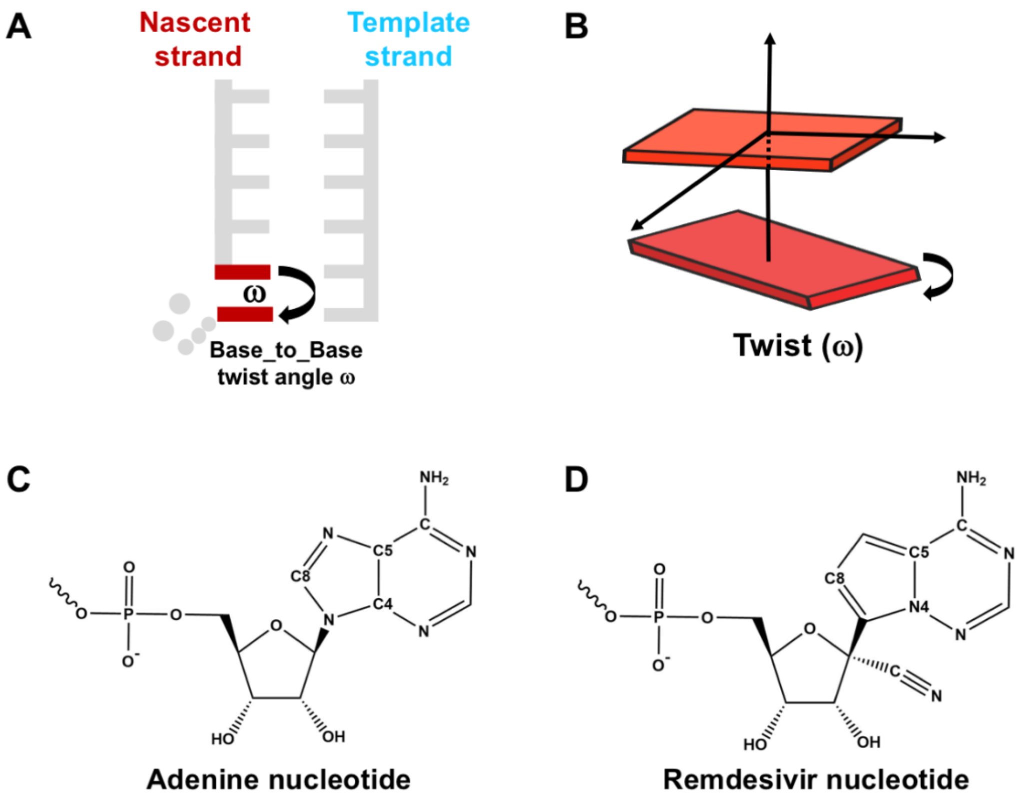


**Figure S3.** Illustration of the twist angle between the base of 3'-terminal nucleotide and the base of ATP/RDV-TP. (**A**) The twist angle was calculated between the 3'-terminal nucleotide and the nucleoside triphosphate. (**B**) Cartoon model illustrates the twist angle. (**C**) The atoms "C4", "C5" and "C8" of adenine nucleotide were used for calculating the twist angle. (**D**) The atoms "N4", "C5" and "C8" of RDV nucleotide were used for calculating the twist angle. See SI Section 4.2 for details.

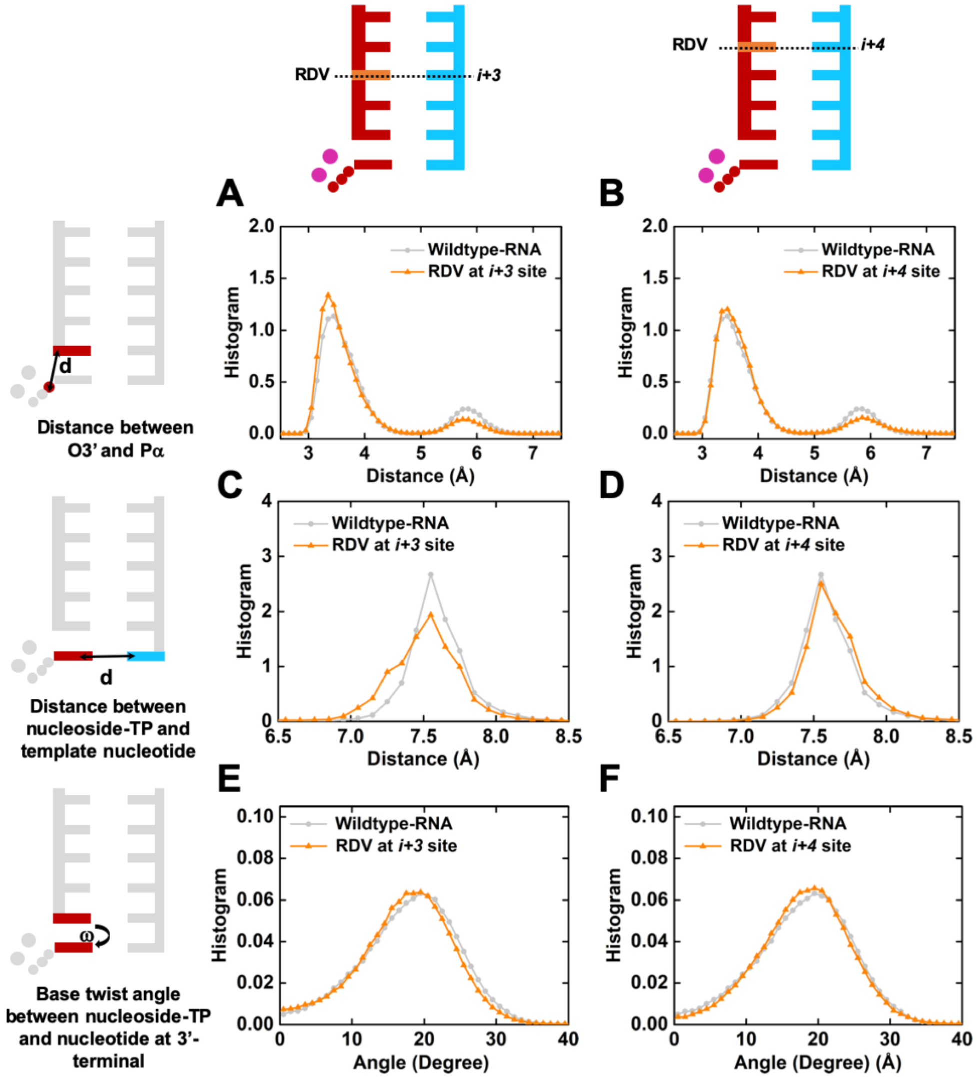


**Figure S4.** Investigation of nucleotide addition in RdRp when RDV is at *i+3* or *i+4* site. The two cartoons in the top horizontal panel denote the site where RDV is positioned (in orange). The three cartoons in the left vertical panel describe the interactions under investigation. For clarification, only the molecules involved in the calculations are colored. Each of the six histograms is calculated for the corresponding structural features (cartoon in the left vertical panel) using the model with RDV located a specific site (cartoon in top horizontal panel). (**A**)-(**B**) Histogram of distance between the Pα of ATP and the O3' atom of the 3'-terminal nucleotide when RDV is at *i+3* (**A**) or *i+4* (**B**) site. (**C**)-(**D**) Histogram of distance between the base of ATP and the base of template nucleotide at *i* site when RDV is at *i+3* (**C**) or *i+4* (**D**) site (see SI Section 4.1 for the details about the distance calculations). (**E**)-(**F**) Histogram of twist angle between the base of ATP and the base of 3'-terminal nucleotide when RDV is at *i+3* (**E**) or *i+4* (**F**) site. See SI Section 4.2 and Supplementary Figure S3 for details about the twist angle calculations in (**E**)-(**F**). In each panel, the histogram for wildtype-RNA is shown in light grey as a reference.

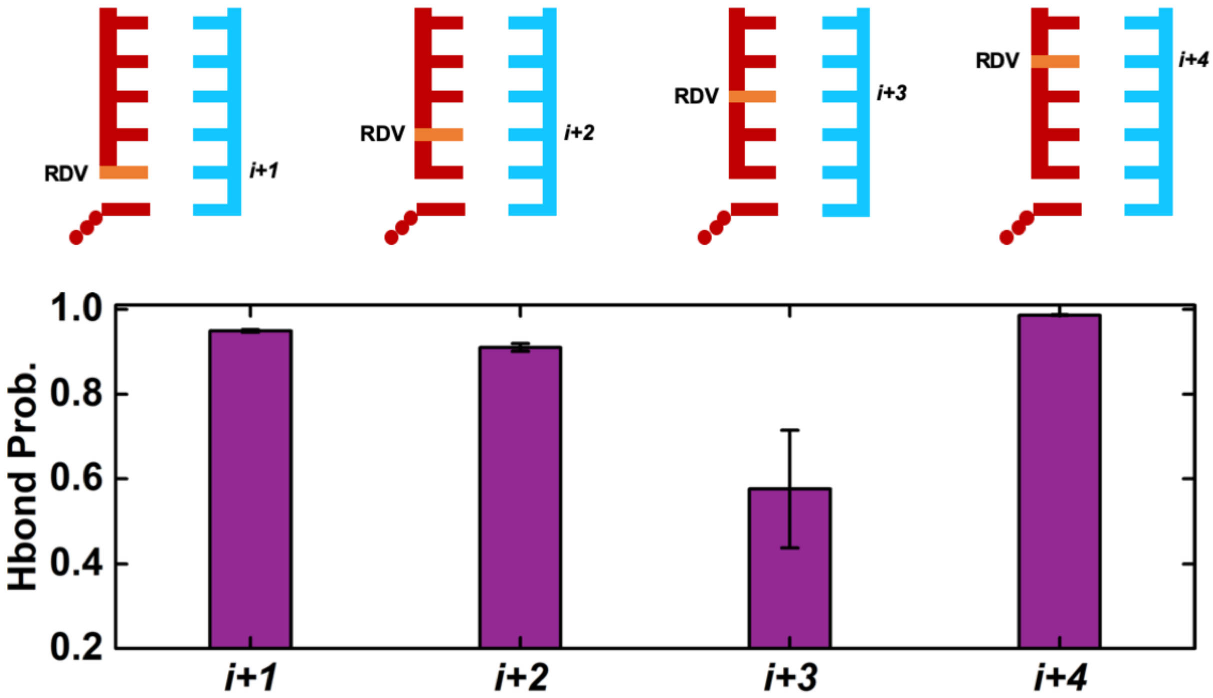


**Figure S5.** Hydrogen bond probability for RDV:U pair at the post-T state with RDV at *i+1*, *i+2*, *i+3*, or *i+4* site. The top panel is the cartoon model of post-T state with RDV at a specific site (in orange). See SI Section 4.4 for details about the calculation of hydrogen bond probability.

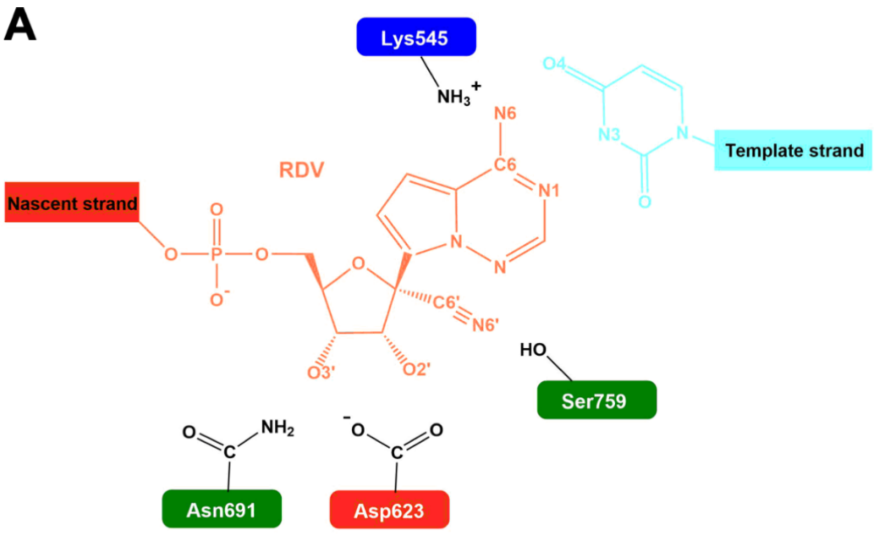


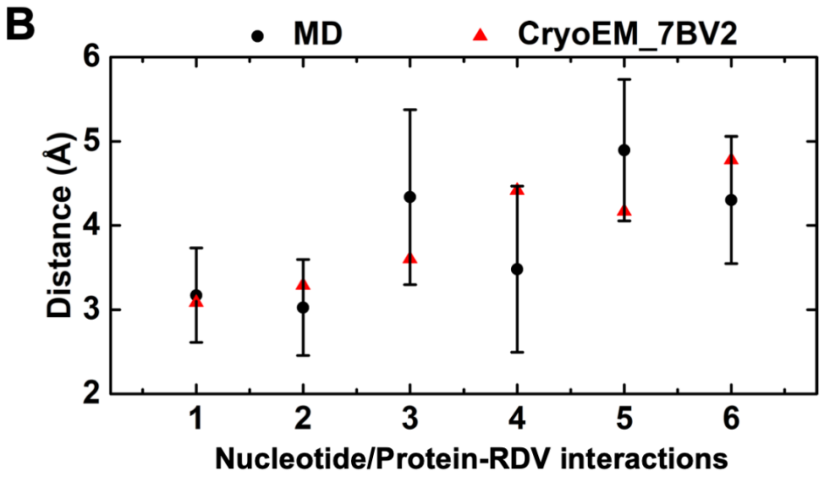


C

| Index | Chain | Site/Resid | Residue | Interacting atom | Chain | Site | Interacting atom |
|---|---|---|---|---|---|---|---|
| 1 | T | *i* | U | N3 | P | *i* | N1 |
| 2 | T | *i* | U | O4 | P | *i* | N6 |
| 3 | A | 691 | Asn | ND2 | P | *i* | O3' |
| 4 | A | 623 | Asp | OD1/OD2 | P | *i* | O3’ |
| 5 | A | 759 | Ser | OG | P | *i* | N6' |
| 6 | A | 545 | Lys | NZ | P | *i* | N6 |

**Figure S6.** Interaction between RDV at *i* site and the surrounding residues in comparison with that of cryo-EM structure (PDBID: 7BV2). (**A**) Schematic representation of RDV at *i* site (orange), surrounded by its base-paired template nucleotide (the base of which is shown in cyan) and protein residues. (**B**) Distances between RDV and its surrounding protein residues/nucleobase calculated using MD conformations (in black circles), in comparison with those calculated from the cryo-EM structure (in red triangles). The means and standard deviations of the distances from MD simulations (after removing the first 10ns from each MD trajectory) were calculated using all the MD conformations of RdRp in the pre-T state with RDV at *i* site. (**C**) Details of the atoms used for the distance calculations.

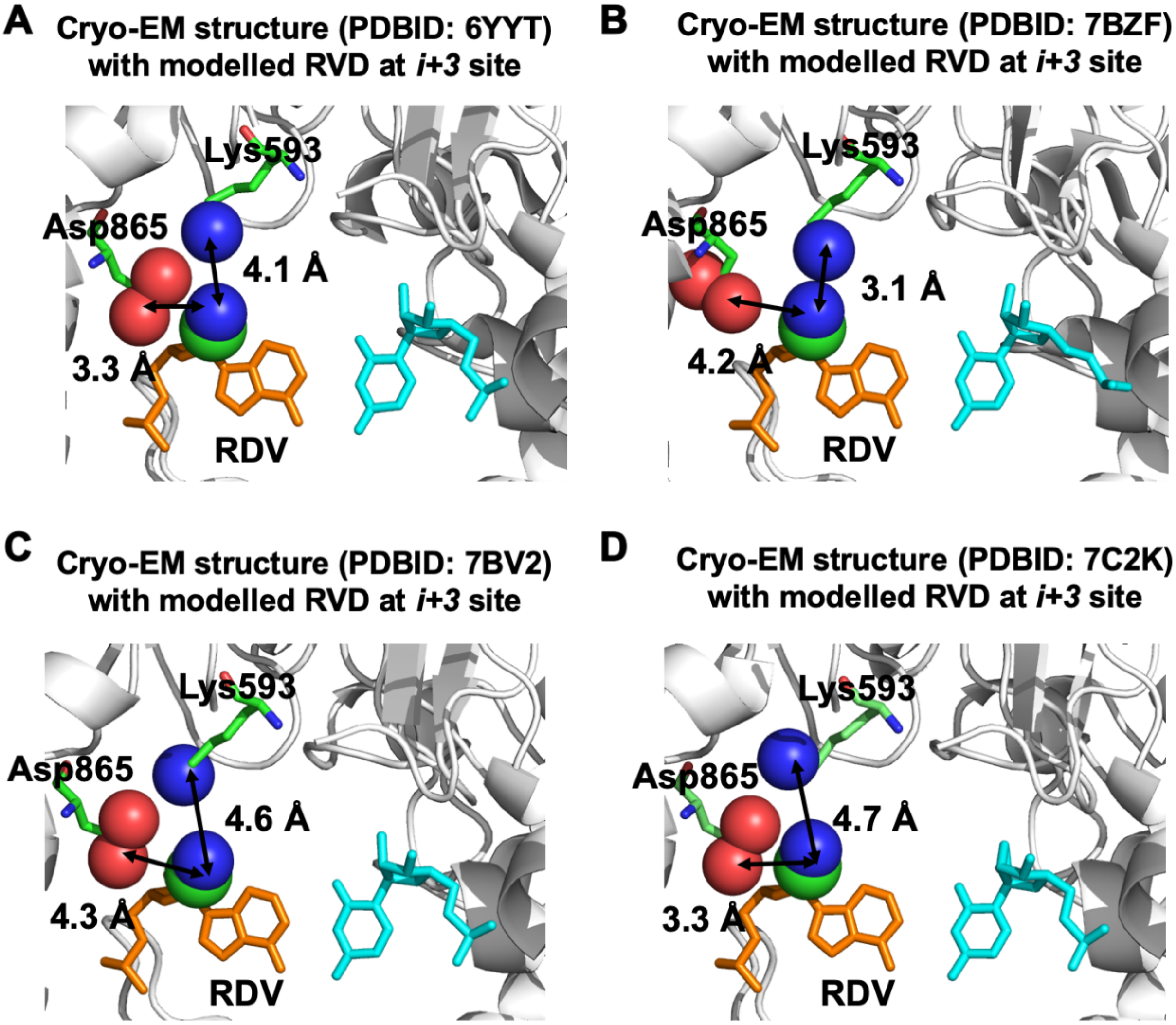


**Figure S7.** RDV at *i+3* site is in a close contact with Lys593 and Asp 865. Cryo-EM structures (**A**-**D** for PDBID: 6YYT, 7BZF, 7BV2 and 7C2K, respectively) with RDV modelled at *i-4* site are shown. The distances between Asp865/Lys593 and the 1'-cyano group are labelled. The "NZ" atom of Lys593 and the nitrogen atom of the 1'-cyano group of RDV are used for calculating the distance between Lys593 and RDV. For the distance between Asp865 and RDV, we measured the minimum distance between the nitrogen atom of 1'-cyano group of RDV and the oxygen atoms in the carbonyl group of Asp865. In each panel, RDV and its template nucleotide is shown in orange and cyan, respectively. The 1'-cyano group of RDV, the oxygens in the carbonyl group of Asp865 and the "NZ" atom of Lys593 are shown in spheres. Protein is displayed in cartoon in the background.

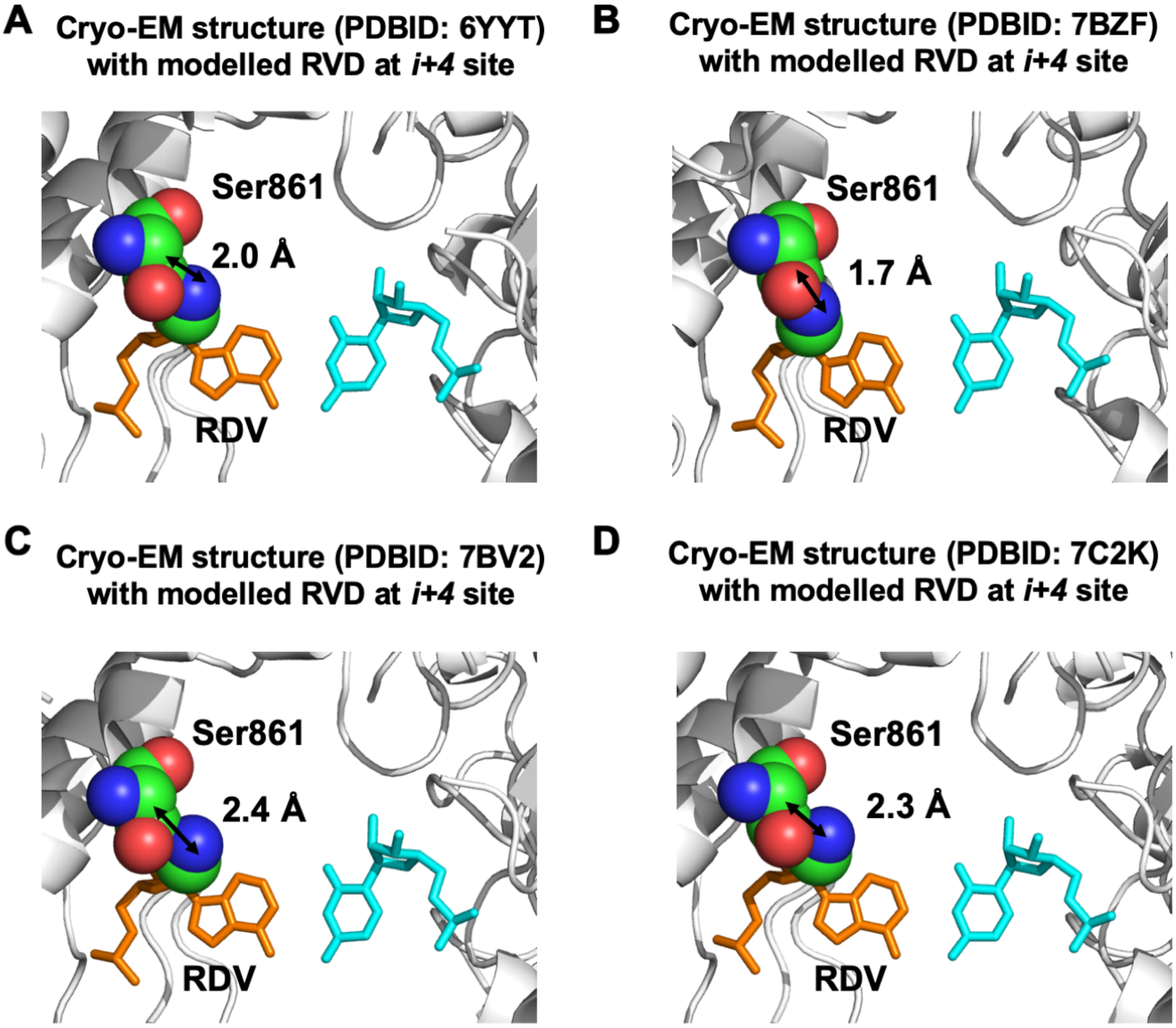


**Figure S8.** Ser861 has steric clash with the 1'-cyano group of RDV at *i+4* site. (**A**) Configuration of Ser861 (shown in spheres) with the 1'-cyano group (shown in spheres) of RDV (in orange) modelled at *i+4* site using the cryo-EM structure (PDBID: 6YYT). The template nucleotide is shown in cyan and protein in shown in cartoon. (**B**)-(**D**) Similar to (A) but for other cryo-EM structures (PDBID: 7BZF in (**B**), 7BV2 in (**C**) and 7C2K in (**D**)). The minimum distance between the heavy atoms of Ser861 and the nitrogen atom in the 1'-cyano group of RDV is labelled in each panel.

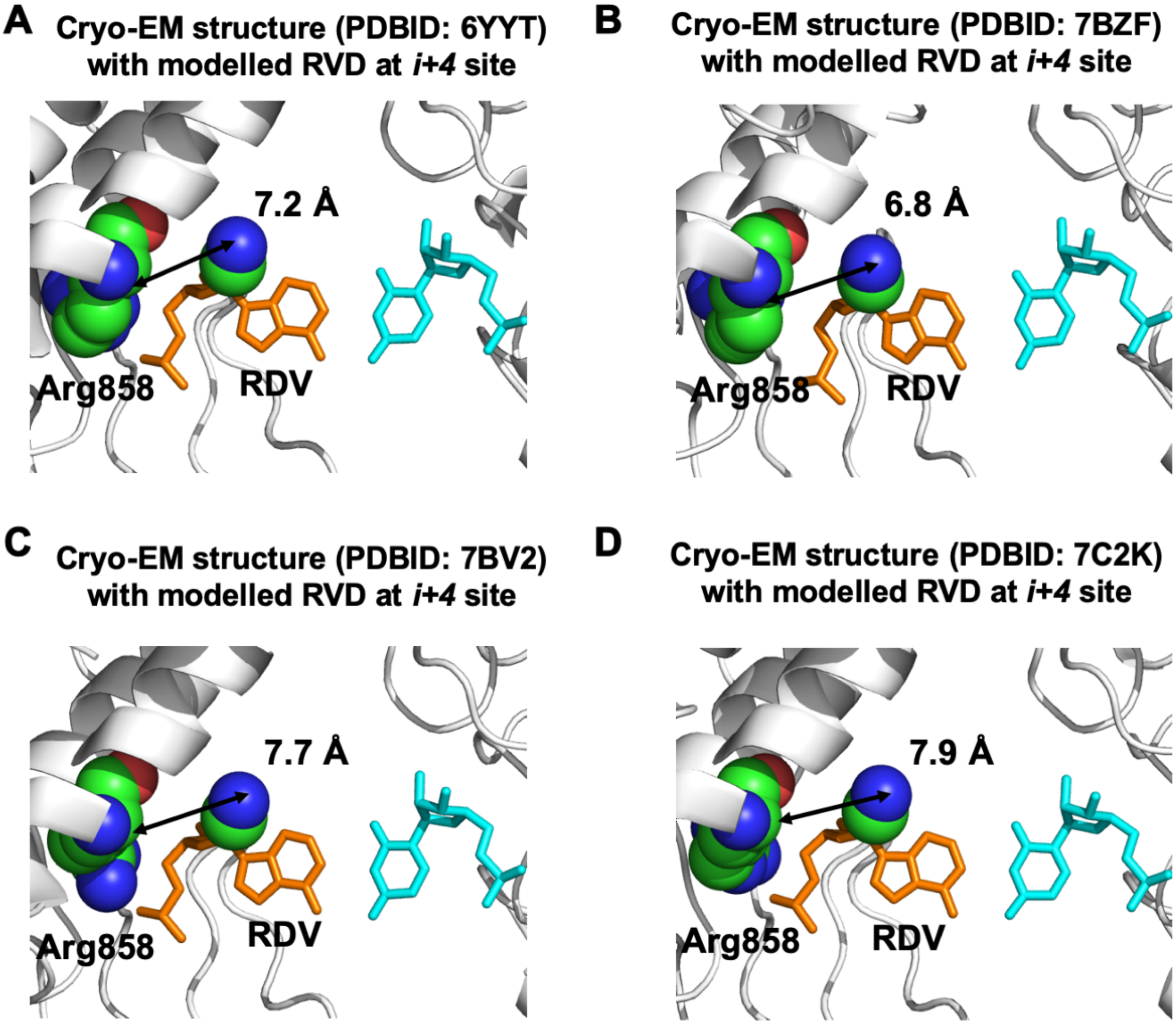


**Figure S9.** Arg858 does not have steric clash with the 1'-cyano group of RDV at *i*+*4* site. (**A**) Cryo-EM structure (PDBID: 6YYT) with RDV modelled at *i*+4 site. Arg858 and the 1'-cyano group of RDV are shown in spheres. RDV and its template nucleotide is shown in orange and cyan, respectively. Protein is displayed in cartoon as the background. (**B**)-(**D**) Similar to (**A**) but for the other three cryo-EM structures (PDBID: 7BZF in (**B**), 7BV2 in (**C**) and 7C2K in (**D**)). The minimum distance between side chain of Arg858 and 1'-cyano group of RDV is labelled in each panel.

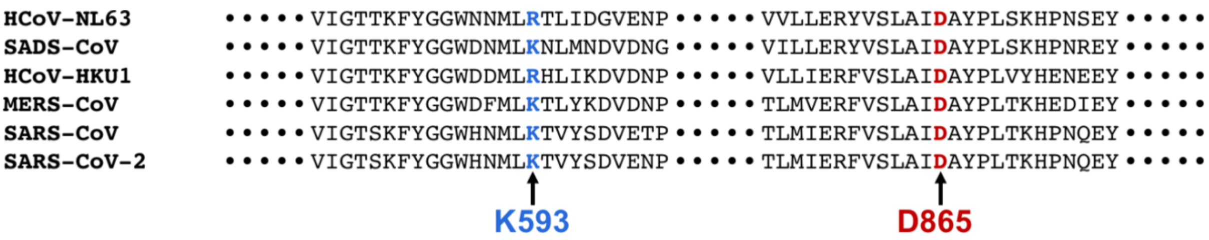


**Figure S10.** Sequence alignment shows the salt bridge $D865^{-}$-$K593^{+}$ is conserved among different coronaviruses, excepting that Lys (K) is replaced with Arg (R) in two human coronaviruses. See SI Section 5 for details about the sequence alignment.

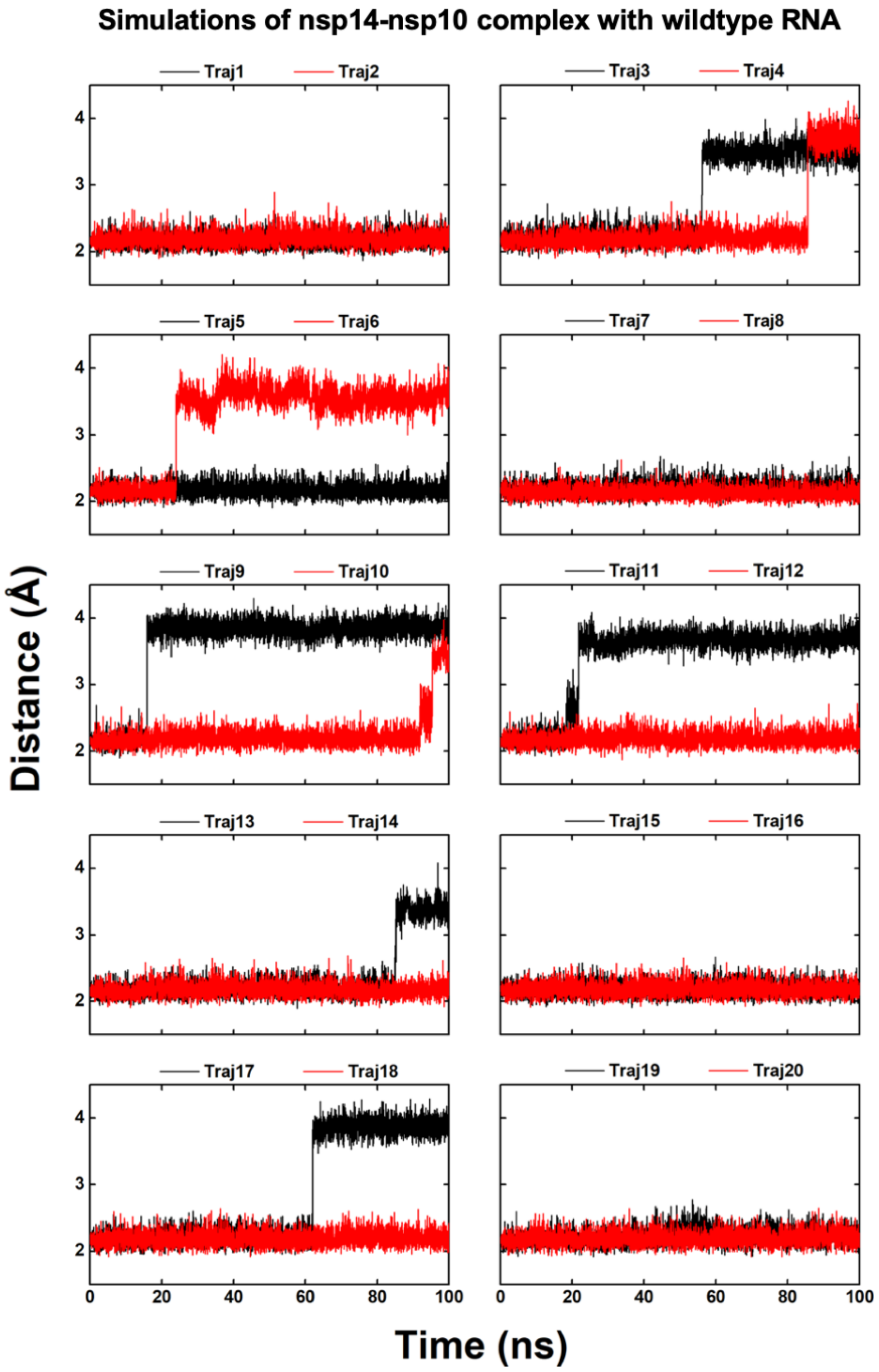


**Figure S11.** The O3'-MgA distance over time for 20 replicas of 100 ns MD simulations of nsp14-nsp10 complex containing wildtype-RNA.

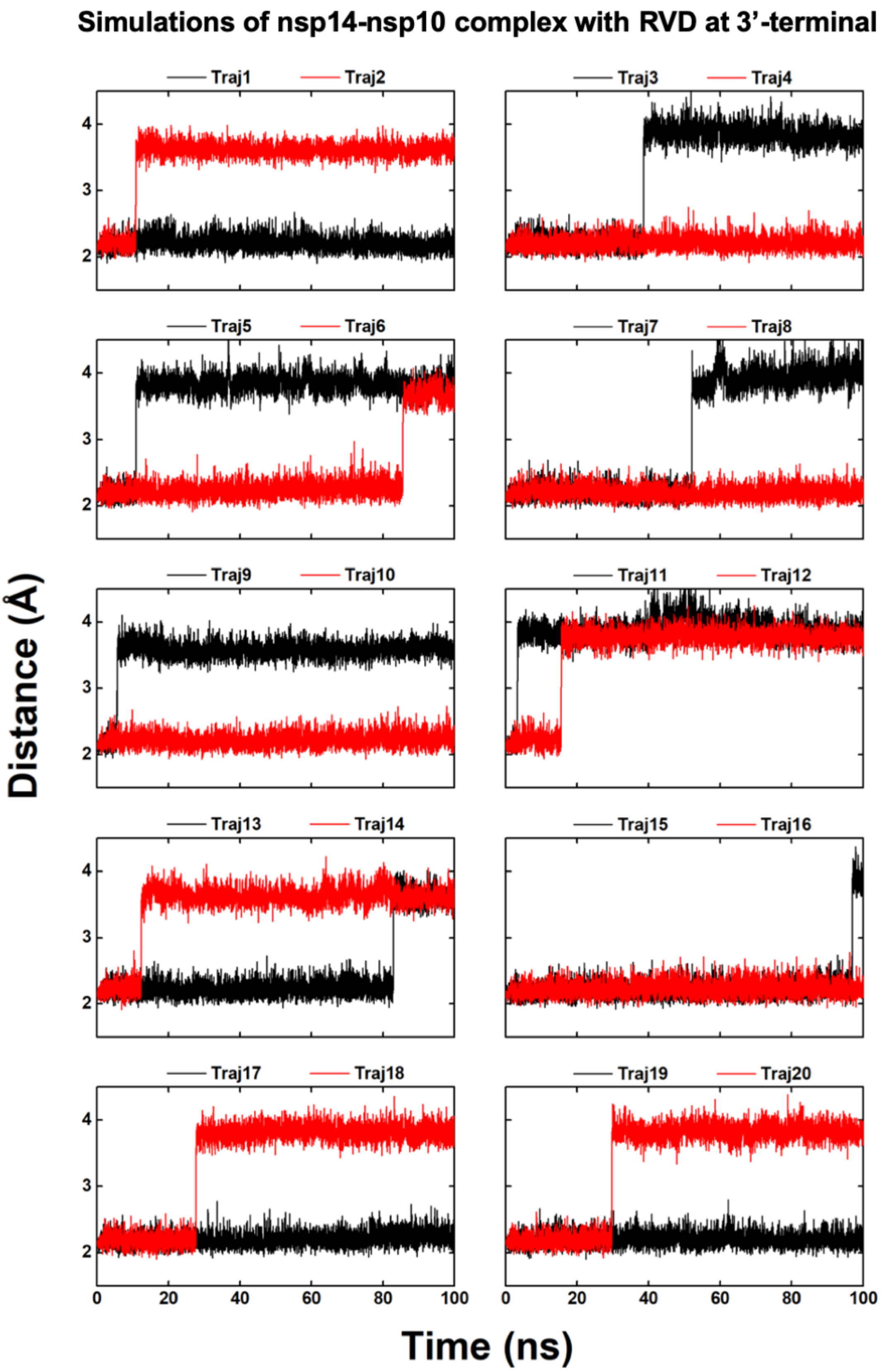


**Figure S12.** The O3’-MgA distance over time for 20 replicas of 100 ns MD simulations of nsp14-nsp10 complex containing single-stranded RNA with RDV at 3’-terminal.

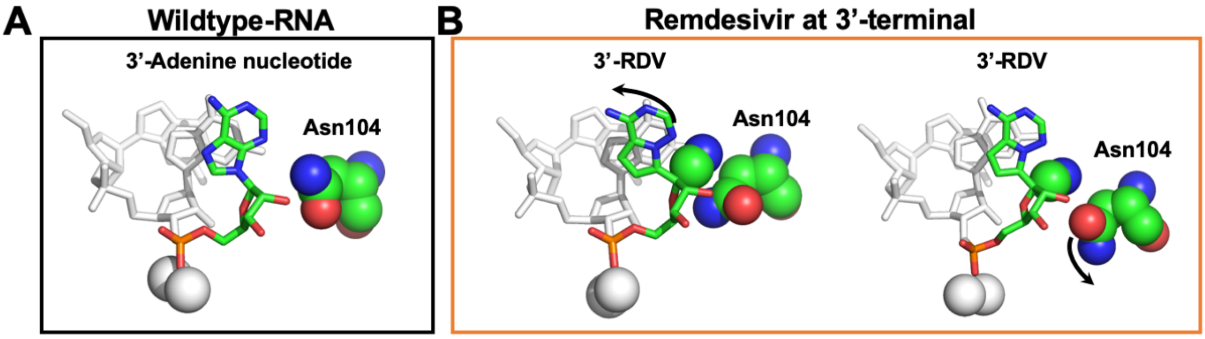


**Figure S13.** Representative conformation of the cleavage site in ExoN. (**A**) Typical MD conformation for wildtype RNA with adenine nucleotide at the 3'-terminal. (**B**) Typical conformations with RDV at 3'-terminal. The 1'-cyano group of RDV and Asn104 are shown in sphere. In (**A**) and (**B**), K-center clustering was performed to divide the MD conformational ensemble into 20 clusters. Only the center conformations of the clusters with population > 20% are shown (see SI Section 4.9 for details).

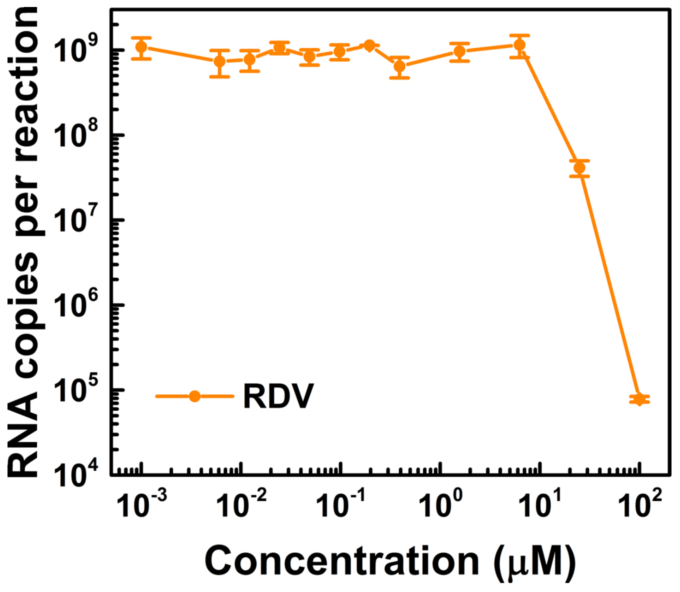


**Figure S14.** Experimental results for the viral RNA copy number under increasing concentration of Remdesivir *in vitro* using live SARS-CoV-2 virus infecting Vero E6 cells. The figure is adapted from our previous work (1).

***Supplementary Text:***

### 1. Structural modeling

***1.1 nsp12-nsp7-nsp8 complex*** The cryo-EM structure of nsp12-nsp7-nsp8 complex (PDBID: 6NUR) (2) was used as the basis to construct the RdRp of SARS-CoV-2. The nearly identical amino acid sequences of nsp12-nsp7-nsp8 complex between SARS-CoV and SARS-CoV-2 (~97.1% sequence identity) render our modeling highly feasible. First, after filling in the missing residues (IDs from 897 to 906) in nsp12, we modelled the nsp12, nsp7 and nsp8 of SARS-CoV-2 based on the corresponding protein subunits of SARS-CoV by modeller9.21 (3). For the homology modelling of nsp12, nsp7 or nsp8, we generated 20 modelled structures and selected the one with the optimal Discrete Optimized Protein Energy (DOPE) assessment score (4) as our final model. Second, the double stranded RNA (dsRNA), ATP and $Mg^{2+}$ ions in the active site were modelled by structural alignment to the norovirus RdRp (PDBID: 3H5Y (5)) using Pymol (6). To facilitate the modelling of Remdesivir at *i* or *i+1* site, we used Coot10.13 (7) to mutate the nucleotides in the corresponding sites to ATP:U and A:U base pairs, respectively. Third, after the structural alignment between SARS-CoV-2 and norovirus (5) RdRps, we found Ser682 of SARS-CoV-2 RdRp is homologous to the Ser300 of norovirus RdRp, but their side chains are at different orientations. To maintain interactions between the side chain of Ser682 and the O2' atom of ATP, we replaced the coordinates of Ser682 with those of Ser300 from the aligned norovirus RdRp (5). Fourth, the protonation states of histidine residues were predicted using propka3.0 module (8) in the pdbpqr2.2.1 (9) package, followed by manual inspection to ensure that the coordination between the $Zn^{2+}$ ion and the corresponding histidine residues (residue IDs 695 and 242 in nsp12) were maintained. Accordingly, histidine with residue IDs 295, 309, 642, 872, and 892 in nsp12, as well as histidine with residue ID 36 in nsp7, have $N_\varepsilon$ atom protonated; the remaining histidine residues have $N_\delta$ atom protonated. The whole complex was placed in a dodecahedron box with the box edges at least 12 Å away from the complex surface. The box was filled with TIP3P water molecules (10), and sufficient counter ions were added to neutralize the whole system. This nsp12-nsp7-nsp8 model containing wildtype RNA with ATP in the active site (*i* site) serves as the structural basis to model Remdesivir in RdRp.

***1.2 nsp14-nsp10 complex*** The crystal structure of nsp14-nsp10 complex of SARS-CoV (PDBID: 5C8S (11)) serves as our modelling template to build the nsp14-nsp10 complex of SARS-CoV-2. First, due to the ~95.7% sequence identity of nsp14-nsp10 complex between

SARS-CoV and SARS-CoV-2, we constructed the model of nsp14-nsp10 complex of SARS-CoV-2 directly by using Pymol (6) to generate the mutations based on the crystal structure of SARS-CoV (11) and ensuring that the side chains of mutated residues could maintain the original orientations. Second, we used modeller9.21 (3) to fill in the missing amino acids (residue IDs 454-464) in nsp14. Third, the ExoN domain of SARS-CoV shares a similar architecture as the proofreading domain of DNA polymerase I Klenow fragment (PDBID: 1KLN (12)) and the ε-subunit of DNA polymerase III (PDBID: 1J53 (13)). Hence, we modelled the single-stranded RNA and $Mg^{2+}$ ions in the catalytic cleavage site by aligning the protein residues in the proofreading domains of these proteins. Specifically, D90, E92, E191, D273, and H268 in nsp14 of SARS-CoV-2, D12, E14, D103, D167 and H162 in the ε-subunit of DNA polymerase III, and E357, D424, D501 and Y497 of domain of DNA polymerase I were used for the structural alignment. The modelled single-stranded RNA contains three nucleotides, because three base pairs are required to be melted to allow the 3'-terminal of nascent strand to access the cleavage site (12). In particular, the 3'-terminal nucleotide and two $Mg^{2+}$ ions were modelled based on the structural alignment to the ε-subunit of DNA polymerase III (13), while the remaining two nucleotides were modelled by aligning to the proofreading domain of DNA polymerase I Klenow fragment (12). Fourth, the alignment between ExoN domains of SARS-CoV-2 and ε-subunit of DNA polymerase III (13) indicate the protein residues (D90, E92, D273 and H268) in SARS-CoV-2 are homologous to the D12, E14, D167 and H162 in DNA polymerase III. To maintain the coordination between these residues and the $Mg^{2+}$ ions in the cleavage site, we replaced the coordinates of D90, E92, D273 and H268 with those of their homologous residues in the aligned ε-subunit of DNA polymerase III (13). We also extracted water molecules coordinated with the $Mg^{2+}$ ions from the aligned ε-subunit of DNA polymerase III (13) to maintain their coordination with the $Mg^{2+}$ ions. Fifth, to facilitate the modelling Remdesivir (adenine nucleoside analogue) at the 3'-terminal, we used Coot10.13 (7) to mutate the 3'-terminal nucleotide to adenine nucleotide. Sixth, protonation states of histidine residues were predicted using the procedure illustrated in Section 1.1 for nsp12-nsp8-nsp7 complex. Specifically, residues 26, 264, 268, 330, 373, 424, 486 and 487 in nsp14 were protonated at $N_{\varepsilon}$, while other histidine residues were protonated at $N_{\delta}$. We did not include the methyl donor and acceptor in our current model, because their binding to the MTase domain of nsp14 does not show obvious structural variations (11). The whole complex was solvated with TIP3P water molecules (10) in a dodecahedron box. The box edges are at least 12 Å away from the complex surface. Enough counter ions were added to neutralize the whole system. This nsp14-nsp10 model has wildtype RNA with adenine nucleotide at the 3'-terminal in ExoN domain and serves the structural basis to model the NTP analogues in ExoN.

### 2. Molecular Dynamics Simulation Set-up

***2.1 Force field parameters*** We used amber99sb-ildn force field (14) to simulate protein and RNA. To generate force field parameters of Remdesivir in either nucleotide or nucleoside triphosphate form, we followed a similar scheme as used in amber99sb-ildn force field (14). In particular, the initial structures for generating force field parameters for Remdesivir were built by modifying the natural adenine nucleotide from 3'-terminal of nascent strand in nsp12 to Remdesivir in the respective form. When deriving the partial charges for Remdesivir, both the 3'- and 5'-terminals of Remdesivir were truncated by hydroxyl groups (as in the nucleoside form). Geometry optimization was performed using HF/6-31G*, followed by single-point calculation with the same method and basis set in Gaussian16 (15). The obtained electrostatic potential was used to generate the partial charges using the Restricted Electrostatic Potential approach (16,17). To make the partial charges of Remdesivir compatible with the existing parameters of RNA molecules, the partial charges of 5'-terminal and 3'-terminal hydroxyl groups were assigned with the empirical values (H5T=0.4295, O5'= −0.6223, H3T=0.4376, O3'= −0.6541) as used in amber99sb-ildn force field (14), and kept constant in the charge fitting. Bonded and Lennard-Jones parameters for Remdesivir were derived from those used for adenine in the amber99sb-ildn force field (14), except those with obvious discrepancy from adenine were instead from General Amber Force Field (18,19). They include the 1'-cyano group as well as the five-membered ring for Remdesivir. For ATP or Remdesivir in the triphosphate form, parameters for the triphosphate tail were taken from those developed by Meagher *et al.* (20).

***2.2 MD simulations for nsp12-nsp7-nsp8 complex***

**2.2.1 MD simulations for nsp12-nsp7-nsp8 complex containing wildtype RNA with ATP in the active site (*i*) site**

We performed multiple steps of energy minimization and position restraint simulations to gradually relax and fully equilibrate the simulation complex containing wildtype RNA with ATP in the active site as follows:

(a) One 10,000-steps energy minimization on the whole system by position restraining the heavy atoms of nucleotides and $Mg^{2+}$ ions with a force constant of 10 kJ×mol$^{-1}$×Å$^{-2}$;

(b) Another 10,000-steps energy minimization without restrain;

(c) One 200 ps position restraint simulation under NVT ensemble (T=300K) with a force constant of 10 kJ×mol$^{-1}$×Å$^{-2}$ on all the heavy atoms of the complex;

(d) Another 500 ps position restraint simulation under NPT ensemble (T = 300 K, P=1 bar) with a force constant of 10 kJ×mol$^{-1}$×Å$^{-2}$ on all the heavy atoms of the complex;

(e) One 10 ns simulation under NPT ensemble (T = 298 K, P = 1 bar) by releasing the position restrain on protein while retaining the position restrain on nucleotides and $Mg^{2+}$ ions;

(f) Another 10 ns simulation under NPT ensemble (T = 298 K, P = 1 bar) was performed without position restraint;

(g) One 100 ns simulation under NPT ensemble (T = 298 K, P = 1 bar) to fully equilibrate the whole system at T = 298 K and P = 1 bar.

It's worthy to note that the structural alignment between SARS-CoV-2 RdRp and norovirus RdRp (5) enables us to pinpoint the residues that may be important for stabilizing the active site. Therefore, we maintained these interactions by adding harmonic constraints in steps (a)-(f). These interactions include: (i) MgA with its coordinated Asp618, Asp760, Asp761 and one $P_\alpha$ oxygen of ATP; (ii) MgB with Asp618, Tyr619, Asp760 and three oxygen atoms from $P_\alpha$, $P_\beta$ and $P_\chi$ of ATP; (iii) hydrogen bond between the 2'-hydroxyl group of ATP and Asn691 (2,5); (iv) hydrogen bond between the 2'-hydroxyl group of ATP and Thr680 (2); (v) hydrogen bond between the 3'-hydroxyl group of ATP and Asp623 (2,5). The harmonic constraints were removed in step (g) for full equilibration of the whole system.

The last configuration of the 100 ns simulation was used to seed 20 replicas of 100 ns MD simulations under NVT ensemble, with simulated annealing from 50 K to 300 K in the first 2 ns. In the 20×100 ns simulations, we added the harmonic constraint to maintain the coordination between the $Zn^{2+}$ ions with their coordinated protein residues ($Zn^{2+}$ ions are located distantly (> 25 Å) from the ATP in the active site): one $Zn^{2+}$ ion is coordinated by His295, Cys301, Cys306 and Cys310; the other one is coordinated by Cys487, His642, Cys645 and Cys646. This is to avoid the $Zn^{2+}$ ions, especially the ones on the surface, from diffusing into the solvent. We applied V-rescale thermostat (21) with the coupling time constant of 0.1 ps. The long-range electrostatic interactions beyond the cut-off at 12 Å were treated with the Particle-Mesh Ewald method (22). Lennard-Jones interactions were smoothly switched off from 10 Å to 12 Å. The neighbour list was updated every 10 steps. An integration time step of 2.0 fs was used and the LINCS algorithm (23) was applied to constrain all bonds. We saved the snapshots every 20 ps, and conformations after 10 ns were collected for subsequent structural analyses. All simulations were performed with Gromacs 5.0 (24).

**2.2.2 MD simulations for nsp12-nsp7-nsp8 complex containing Remdesivir**

We modelled Remdesivir in the nucleotide form at the corresponding site of the nascent strand based on the last configuration of the 100 ns equilibrated simulation containing wildtype RNA with ATP in the active site (step (g) in Section 2.2.1):

- Modeling Remdesivir at *i* site: we replaced ATP with Remdesivir in the triphosphate form at the active site (*i* site);
- Modeling Remdesivir at *i+1* site: we substituted the 3'-adenine nucleotide with Remdesivir;
- Modeling the post-translocation (post-T) state with Remdesivir at *i+2*, *i+3* or *i+4* site: we replaced the nucleotide in the corresponding site with Remdesivir;
- Modeling the pre-translocation (pre-T) state with Remdesivir at *i*, *i+1*, *i+2 or i+3* site: we added covalent bond between the O3' atom of 3'-adenine nucleotide and the Pα atom of ATP, and replaced the triphosphate tail with hydrogen atom. We removed MgB from the model. The nucleotide at corresponding upstream site of the nascent strand was substituted with Remdesivir.

For each model with Remdesivir, the whole system was relaxed gradually for the equilibration:

(a) One 10,000-steps energy minimization with position restraint (10 $kJ \times mol^{-1} \times Å^{-2}$) on the heavy atoms of nucleotides and $Mg^{2+}$ ions;
(b) Another 10,000-steps energy minimization without position restraint;
(c) The energy minimized system was further relaxed by 200 ps simulation under NVT ensemble (T=300 K) with position restrain (10 $kJ \times mol^{-1} \times Å^{-2}$) on all the heavy atoms of the system;
(d) Another 500 ps simulation under NPT ensemble (T=300 K, P=1 bar) with position restrain (10 $kJ \times mol^{-1} \times Å^{-2}$) on all the heavy atoms of the system.

The last configuration of the position restraint simulation was utilized to run 20 replicas of 100 ns NVT simulations (T=300 K) with different random seeds. Temperature was gradually increased from 50 K to 300 K in the first 2 ns using simulated annealing. In the 20×100 ns simulations, harmonic constraint was added between the $Zn^{2+}$ ions and their coordinated protein residues to maintain their coordination (one $Zn^{2+}$ ion with His295, Cys301, Cys306 and Cys310; the other $Zn^{2+}$ ion with Cys487, His642, Cys645 and Cys646). Same parameters as used in Section 2.2.1 were utilized for MD simulations. MD snapshot was saved every 20 ps and conformations after 10 ns were collected for subsequent structural analyses.

**2.3** ***MD simulations for nsp14-nsp10 complex***

#### 2.3.1 MD simulations for nsp14-nsp10 complex containing wildtype RNA with adenine nucleotide at the 3'-terminal of nascent strand

We performed multiple steps of energy minimization and position restraint simulations to gradually relax and fully equilibrate the simulation complex as follows:

(a) One 10,000-steps energy minimization with position restraint (10 $kJ\times mol^{-1}\times Å^{-2}$) on the heavy atoms of nucleotides and $Mg^{2+}$ ions;
(b) Another 10,000-steps energy minimization on the whole system without position restraint;
(c) One position restraint simulation of 200 ps under NVT ensemble (T=300 K) by using a restraint force constant of 10 $kJ\times mol^{-1}\times Å^{-2}$ on all the heavy atoms of the nsp14-nsp10 complex;
(d) Another 500 ps NPT ensemble (T=300 K, P=1 bar) with position restraint (10 $kJ\times mol^{-1}\times Å^{-2}$) on all the heavy atoms of the nsp14-nsp10 complex;
(e) One 10 ns NPT simulation (T=300 K, P=1 bar) with position restraint on the nucleotides (10 $kJ\times mol^{-1}\times Å^{-2}$) and $Mg^{2+}$ ions (100 $kJ\times mol^{-1}\times Å^{-2}$). In this step, we added a stronger restraining force constant (100 $kJ\times mol^{-1}\times Å^{-2}$) on $Mg^{2+}$ ions, in order to reduce the perturbation caused by the full relaxation of the surrounding protein residues;
(f) The position restraint on nucleotides was released and the system was simulated for another 30 ns under NVT ensemble (T=300 K) for full equilibration. Simulated annealing was performed from 50 K to 300 K in the first 2 ns.

We have added harmonic constraint between $Mg^{2+}$ ions and their coordinated residues in steps (a)-(d). The coordination network of two $Mg^{2+}$ ions includes: MgA with carboxyl group of Asp90, oxygen atom in the phosphate backbone of 3'-adenine nucleotide and the O3' atom of the guanine nucleotide next to the 3'-adenine nucleotide; MgB with carboxyl group of Asp90, carboxyl group of Asp273, carboxyl group of Glu92 and oxygen atom in the phosphate backbone of 3'-adenine nucleotide. The constraint has been removed after step (d).

The last configuration of the 30 ns simulation was used to seed 20 replicas of 100 ns NVT simulations (T=300 K), including the simulated annealing from 50 K to 300 K in the first 2 ns. It is worthy to note that the nascent RNA strand will backtrack from nsp12 and protrude into the cleavage site of nsp14 for excision. However, there is no available structural information for the binding interfaces between nsp12 and nsp14. Therefore, in order to recapitulate the condition in which the backtracked RNA enters the active site of nsp14 through the binding interface between nsp12 and nsp14, we placed a position restraint (100 $kJ \times mol^{-1} \times Å^{-2}$) on the 5'-nucleotide and the phosphate atom of the neighboring nucleotide to mimic the conformational constrain imposed by the upstream RNA strand. In addition, we also added a weak position restraint (10 $kJ \times mol^{-1} \times Å^{-2}$) on the $Mg^{2+}$ ions to maintain their coordination. Furthermore, harmonic constraint was added between the $Zn^{2+}$ ions and their coordinated protein residues to maintain their coordination ($Zn^{2+}$ ions are located distantly (> 14 Å) from nucleotides in the cleavage site). The coordination network of the five $Zn^{2+}$ ions includes: (i) $Zn^{2+}$ ion (residue ID 201 in nsp10) with Cys74, Cys77, His83 and Cys90; (ii) $Zn^{2+}$ ion (residue ID 202 in nsp10) with Cys117, Cys120, Cys128 and Cys130; (iii) $Zn^{2+}$ ion (residue ID 601 in nsp14) with Cys207, Cys210, Cys226 and His229; (iv) $Zn^{2+}$ ion (residue ID 602 in nsp14) with His257, Cys261, His264 and Cys279; (v) $Zn^{2+}$ ion (residue ID 603 in nsp14) with Cys452, Cys477, Cys484 and His487. The same parameters as used above for the RdRp production simulations (Section 2.2.1) were applied. The MD snapshot was saved every 20 ps and conformations after 10 ns were collected for subsequent structural analyses.

#### 2.3.2 MD simulations for nsp14-nsp10 complex containing Remdesivir at the 3'-terminal of nascent strand

We constructed the system containing Remdesivir in the nucleotide form at the 3'-terminal of nascent strand based on the last configuration of the 30 ns equilibrated simulation containing 3'-adenine nucleotide. Specifically, the 3'-adenine nucleotide was replaced with Remdesivir.

For each system, we performed multiple steps of energy minimization and position restraint simulations for full equilibration as follows:

(a) One 10,000-steps energy minimization with position restraint (10 $kJ \times mol^{-1} \times Å^{-2}$) on the heavy atoms of nucleotides and $Mg^{2+}$ ions;

(b) Another 10,000-steps energy minimization on the whole system without position restraint;

(c) One 200 ps NVT (T=300 K) position restraint simulation with a force constant of 10 $kJ \times mol^{-1} \times Å^{-2}$ on all the heavy atoms of the complex;

(d) Another 500 ps NPT (T=300 K, P=1 bar) position restraint simulation with a force constant of 10 $kJ \times mol^{-1} \times Å^{-2}$ on all the heavy atoms of the complex.

The last configuration of the position restraint simulations was used to randomly seed 20 replicas of 100 ns MD simulations under NVT ensemble (T=300 K), with simulated annealing from 50 K to 300 K in the first 2 ns. Harmonic constraint was added to maintain the coordination between $Zn^{2+}$ ions and their coordinated protein residues, as described in the Section 2.3.1. Position restraint was applied on the 5'-terminal nucleotides and $Mg^{2+}$ ions in the same way as used in the production simulation in Section 2.3.1. The MD snapshot was saved every 20 ps and conformations after 10 ns were collected for subsequent structural analyses. Same parameters as used above for the RdRp simulations (Section 2.2.1) were utilized.

### 3. Generation of translocation pathway

To investigate if the translocation of Remdesivir from *i+3* to *i+4* site is inhibited, we used the Climber algorithm (25,26) to generate a translocation pathway.

***3.1 Modelling the pre-T and post-T states of SARS-CoV2-RdRp.*** First, we constructed the pre-T conformation based on the configuration of pre-T state with Remdesivir at *i+3* site after position restraint simulations (Section 2.2.2) and replaced Remdesivir with adenine nucleotide. Second, we constructed post-T conformation based on the configuration of post-T state with Remdesivir at *i+4* site after position restraint simulations (Section 2.2.2) and replaced Remdesivir with adenine nucleotide. Third, in the translocation, all the nucleotides are translocating forward by one base pair position. Our structural model of SARS-CoV-2 RdRp contains only the coordinates of nucleotides from *i-1* to *i+7* site, which enables us to simulate the translocation of nucleotides from *i-1* to *i+6* site in the pre-T conformation to *i* to *i+7* site in the post-T conformation. Therefore, we removed the most upstream base pair of nucleotides at *i+7* site in the pre-T conformation, as well as the ATP and the most downstream nucleotide at *i-1* site in the post-T conformation. Nucleotides were mutated by Coot10.13 (7) to make the nucleotides' sequence consistent between pre-T and post-T states. Fourth, energy minimization was performed for the pre-T and post-T conformations using the Energy Calculation and Dynamics (ENCAD) simulation program (27). The energy-minimized pre-T and post-T conformations were used as the initial and final conformation for generating the preliminary pathway, respectively.

***3.2 Generation of translocation pathway.*** We used Climber algorithm (25,26) to generate a low-energy pathway for the translocation of double-stranded RNA (dsRNA) containing an adenine nucleotide moving from *i+3* to *i+4* site in SARS-CoV-2 RdRp. An external energy was applied on dsDNA to drive the system from the initial pre-T state to the final post-T state. In particular, this external energy is consisted of a series of harmonic potentials applying on distances between atom pairs (one atom belongs to dsRNA, while the other atom belongs to protein and ions) (25,26). To generate the translocation pathway, we performed a 350-step Climber simulation, which gradually drives the dsRNA from the pre-T state to the post-T state. In this simulation, the system has succeeded in translocation with a RMSD of 1.8 Å to the post-T state. Next, we chose conformations along this translocation pathway every 10 steps and replaced the adenine nucleotide (translocating from *i+3* to *i+4* site) with Remdesivir. Subsequent energy minimization of each of these conformations was then performed by Gromacs 5.0 (24).

### 4. Structural analyses

***4.1 Base-to-base distance for the base pairing at the active site of RdRp*** We used the center of mass (c.o.m.) of the ring directly connected to the ribose to determine the base position of the nucleotides or Remdesivir involved in the distance calculations. Only heavy atoms constituted the ring were considered in the c.o.m. calculation.

***4.2 Twist angle for the base stacking at the active site of RdRp*** Adenine and Remdesivir nucleotides have two conjugated rings in the base, and we used the vector formed by the two atoms in the connected region of the two rings to measure the twist angle (Fig. S3). In adenine, the vectors formed by the "C4" and "C5" atoms were used (Fig. S3C), while in Remdesivir, the "N4" and "C5" were utilized for defining the vector (Fig. S3D). The vector in the base of ATP/Remdesivir-TP was projected onto the plane defined by the three atoms in the base of 3'-terminal nucleotide ("C8", "C5" and "C4" for adenine nucleotide, "C8", "C5" and "C4" for Remdesivir, Figs. S3C and S3D). The angle between the projected vector and the vector in the plane was calculated as the base-to-base twist angle.

***4.3 Root Mean Square Deviation (RMSD) calculations of MD conformations based on our structural model against the cryo-EM structures*** We compared RMSDs of protein residues that are present in both of our structural model and cryo-EM structures (cryo-EM structures contain longer N-terminal tail and nucleotides). In particular, we used the Cα atoms of residues 118 to 919 (excluding residues 896 to 911) for comparing the structural similarity of protein between our structural model and the cryo-EM structures. When comparing the structural similarity of nucleic acids, we only included nucleotides present in both our structural model and the cryo-EM structures. Specifically, nucleotides in the template strand from $i$ to $i+7$ sites and in the nascent strand from $i$ to $i+7$ sites were included, and their phosphate backbones and heavy atoms in the ribose ring were used for the RMSD calculations. When Remdesivir is present (PDBID: 7BV2 (28) with Remdesivir at $i$ site and 7C2K (29) with Remdesivir at $i+1$ site), all the heavy atoms of Remdesivir were included for the RMSD measurement. RMSDs were calculated based on the MD conformational ensemble using bootstrap algorithm. In each bootstrap sample, 20 trajectories were randomly selected with replacement. The mean values and standard deviations were estimated by 20 bootstrap samples.

***4.4 Hydrogen bond probability of base pairs*** Hydrogen bond probability was calculated by the apparent number of hydrogen bonds divided by the optimal number of hydrogen bonds. Bootstrap algorithm was used by generating 20 bootstrap samples to calculate hydrogen bond probability. For each sample, 20 trajectories were randomly selected with replacement.

***4.5 Interaction energy between 1'-cyano group and the side chains of K593 and D865*** We collected the MD conformations of pre-T state with Remdesivir at $i+3$ site to calculate the interaction energy between 1'-cyano group of Remdesivir and the side chains of K593 and D865. Only the conformations with Remdesivir:U in their canonical base paired configuration (two hydrogen bonds formed between Remdesivir and uracil) were considered in the calculations. To set up the calculations, both electrostatic interactions and Lennard-Jones interactions were cut off at 30 Å. Water and counter ions were removed to accelerate the calculations. The side chain of K593, the side chain of D865 and the 1'-cyano group of Remdesivir were treated as three energy groups, and the non-bonded interaction energies among the three energy groups were calculated. Bootstrap method was applied to estimate the means and standard deviations. Specifically, after removing the MD conformations in the first 10 ns, canonical base paired configurations (two hydrogen bonds for the Remdesivir:U pair) were observed in 17 simulations. Thus, we generated 17 bootstrap samples, each of which contains 17 random trajectories selected with replacement. Afterward, the mean values and standard deviations were derived from the 17 samples.

***4.6 Distance between 1'-cyano group and K593/D865/S861 in the translocation of Remdesivir from i+3 to i+4 site*** The nitrogen atom in the 1'-cyano group of Remdesivir was used to calculate its distance to the nitrogen in the quaternary amine group of K593 and the backbone oxygen atom of S861. Because the two oxygen atoms in the carbonyl group of D865 are chemically equivalent, we calculated the minimum distance between the nitrogen atom in the 1'-cyano group of Remdesivir and the two oxygen atoms in the carbonyl group of D865.

***4.7 RMSD to the pre-T state in the translocation of Remdesivir from i+3 to i+4 site*** We aligned the energy minimized conformations along the translocation pathway to the energy minimized pre-T conformation by the Cα atoms of protein and then calculated RMSD of the heavy atoms of nucleotides and Remdesivir.

***4.8 Distance between Asn104 and adenine or NTP analogue at the 3'-terminal site in ExoN*** We used the c.o.m. of the ring directly connected to the ribose to determine the base position of 3'-terminal adenine/Remdesivir nucleotide. For Asn104, the nitrogen atom in the amide group was used for the distance calculations.

***4.9 Selection of representative conformations in ExoN*** We performed K-center clustering (30) based on the RMSD of heavy atoms in single-strand RNA and Asn104. The MD conformations were divided into 20 clusters. Only the center conformation in the cluster with population > 20% is present in Fig. S13. In particular, for the wildtype system with 3'-adenine nucleotide, the representative conformation is in the cluster of population 90%. For the system with 3'-Remdesivir nucleotide, the two representative conformations are in the clusters of population 46% and 26%, respectively.

### 5. Sequence alignment

The sequence of nsp12 SARS-CoV-2 was compared with the GenBank (31) sequences of several CoVs (Fig. S10): Human coronavirus NL63 (HCoV-NL63, accession code YP_003766.2), swine acute diarrhea syndrome coronavirus (SADS-CoV, accession code QID98967.1), Human coronavirus NL63 (HCoV-HKU1, accession code AGW27852.1), Middle East respiratory syndrome-related coronavirus (MERS-CoV, accession code

YP_009047223.1), SARS-CoV (accession code AEA10937.1). The sequence alignment was performed by Clustal Omega1.2.4 (32).

### 6. Reference


1. Choy, K.-T., Wong, A.Y.-L., Kaewpreedee, P., Sia, S.-F., Chen, D., Hui, K.P.Y., Chu, D.K.W., Chan, M.C.W., Cheung, P.P.-H. and Huang, X. (2020) Remdesivir, lopinavir, emetine, and homoharringtonine inhibit SARS-CoV-2 replication *in vitro*. *Antiviral Res.*, 104786.
2. Kirchdoerfer, R.N. and Ward, A.B. (2019) Structure of the SARS-CoV nsp12 polymerase bound to nsp7 and nsp8 co-factors. *Nat. Commun.*, **10**, 2342.
3. Webb, B. and Sali, A. (2016) Comparative protein structure modeling using MODELLER. *Curr. Protoc. Bioinf.*, **15**, 5-6.
4. Shen, M.-y. and Sali, A. (2006) Statistical potential for assessment and prediction of protein structures. *Protein Sci.*, **15**, 2507-2524.
5. Zamyatkin, D.F., Parra, F., Machín, Á., Grochulski, P. and Ng, K.K.S. (2009) Binding of 2′-amino-2′-deoxycytidine-5′-triphosphate to Norovirus polymerase induces rearrangement of the active site. *J. Mol. Biol.*, **390**, 10-16.
6. Imbert, I., Guillemot, J.-C., Bourhis, J.-M., Bussetta, C., Coutard, B., Egloff, M.-P., Ferron, F., Gorbalenya, A.E. and Canard, B. (2006) A second, non-canonical RNA-dependent RNA polymerase in SARS coronavirus. *EMBO J.*, **25**, 4933-4942.
7. Emsley, P., Lohkamp, B., Scott, W.G. and Cowtan, K. (2010) Features and development of Coot. *Acta Crystallogr., Sect. D: Biol. Crystallogr.*, **66**, 486-501.
8. Olsson, M.H.M., Søndergaard, C.R., Rostkowski, M. and Jensen, J.H. (2011) PROPKA3: Consistent treatment of internal and surface residues in empirical pKa predictions. *J. Chem. Theory Comput.*, **7**, 525-537.
9. Dolinsky, T.J., Nielsen, J.E., McCammon, J.A. and Baker, N.A. (2004) PDB2PQR: an automated pipeline for the setup of Poisson–Boltzmann electrostatics calculations. *Nucleic Acids Res.*, **32**, W665-W667.
10. Jorgensen, W.L., Chandrasekhar, J., Madura, J.D., Impey, R.W. and Klein, M.L. (1983) Comparison of simple potential functions for simulating liquid water. *J. Chem. Phys.*, **79**, 926-935.
11. Ma, Y., Wu, L., Shaw, N., Gao, Y., Wang, J., Sun, Y., Lou, Z., Yan, L., Zhang, R. and Rao, Z. (2015) Structural basis and functional analysis of the SARS coronavirus nsp14–nsp10 complex. *Proc. Natl. Acad. Sci. U. S. A.*, **112**, 9436.
12. Beese, L.S., Derbyshire, V. and Steitz, T.A. (1993) Structure of DNA polymerase I Klenow fragment bound to duplex DNA. *Science*, **260**, 352-355.
13. Hamdan, S., Carr, P.D., Brown, S.E., Ollis, D.L. and Dixon, N.E. (2002) Structural basis for proofreading during replication of the *Escherichia coli* chromosome. *Structure*, **10**, 535-546.

14. Lindorff-Larsen, K., Piana, S., Palmo, K., Maragakis, P., Klepeis, J.L., Dror, R.O. and Shaw, D.E. (2010) Improved side-chain torsion potentials for the Amber ff99SB protein force field. *Proteins: Struct., Funct., Bioinf.*, **78**, 1950-1958.
15. Frisch, M.J., Trucks, G.W., Schlegel, H.B., Scuseria, G.E., Robb, M.A., Cheeseman, J.R., Scalmani, G., Barone, V., Petersson, G.A., Nakatsuji, H. *et al.* (2016) Gaussian 16 Rev. C.01. Gaussian, Inc., Wallingford, CT.
16. Woods, R.J. and Chappelle, R. (2000) Restrained electrostatic potential atomic partial charges for condensed-phase simulations of carbohydrates. *J. Mol. Struct.: THEOCHEM*, **527**, 149-156.
17. Wang, J., Cieplak, P. and Kollman, P.A. (2000) How well does a restrained electrostatic potential (RESP) model perform in calculating conformational energies of organic and biological molecules? *J. Comput. Chem.*, **21**, 1049-1074.
18. Wang, J.M., Wolf, R.M., Caldwell, J.W., Kollman, P.A. and Case, D.A. (2004) Development and testing of a general amber force field. *J. Comput. Chem.*, **25**, 1157-1174.
19. Wang, J., Wang, W., Kollman, P.A. and Case, D.A. (2006) Automatic atom type and bond type perception in molecular mechanical calculations. *J. Mol. Graphics Modell.*, **25**, 247-260.
20. Meagher, K.L., Redman, L.T. and Carlson, H.A. (2003) Development of polyphosphate parameters for use with the AMBER force field. *J. Comput. Chem.*, **24**, 1016-1025.
21. Bussi, G., Donadio, D. and Parrinello, M. (2007) Canonical sampling through velocity rescaling. *J. Chem. Phys.*, **126**, 014101.
22. Essmann, U., Perera, L., Berkowitz, M.L., Darden, T., Lee, H. and Pedersen, L.G. (1995) A smooth particle mesh Ewald method. *J. Chem. Phys.*, **103**, 8577-8593.
23. Hess, B., Bekker, H., Berendsen, H.J.C. and Fraaije, J.G.E.M. (1997) LINCS: A linear constraint solver for molecular simulations. *J. Comput. Chem.*, **18**, 1463-1472.
24. Abraham, M.J., Murtola, T., Schulz, R., Páll, S., Smith, J.C., Hess, B. and Lindahl, E. (2015) GROMACS: High performance molecular simulations through multi-level parallelism from laptops to supercomputers. *SoftwareX*, **1**, 19-25.
25. Weiss, D.R. and Levitt, M. (2009) Can morphing methods predict intermediate structures? *J. Mol. Biol.*, **385**, 665-674.
26. Silva, D.A., Weiss, D.R., Avila, F.P., Da, L.T., Levitt, M., Wang, D. and Huang, X.H. (2014) Millisecond dynamics of RNA polymerase II translocation at atomic resolution. *Proc. Natl. Acad. Sci. U. S. A.*, **111**, 7665-7670.
27. Levitt, M. (1983) Molecular-dynamics of native protein .1. Computer-simulation of trajectories. *J. Mol. Biol.*, **168**, 595-620.

28. Yin, W., Mao, C., Luan, X., Shen, D.-D., Shen, Q., Su, H., Wang, X., Zhou, F., Zhao, W., Gao, M. *et al.* (2020) Structural basis for inhibition of the RNA-dependent RNA polymerase from SARS-CoV-2 by Remdesivir. *Science*, **368**, 1499-1504.
29. Wang, Q., Wu, J., Wang, H., Gao, Y., Liu, Q., Mu, A., Ji, W., Yan, L., Zhu, Y., Zhu, C. *et al.* (2020) Structural basis for RNA replication by the SARS-CoV-2 polymerase. *Cell*, **182**, 417-428.
30. Bowman, G.R., Beauchamp, K.A., Boxer, G. and Pande, V.S. (2009) Progress and challenges in the automated construction of Markov State Models for full protein systems. *J. Chem. Phys.*, **131**, 124101.
31. Benson, D.A., Cavanaugh, M., Clark, K., Karsch-Mizrachi, I., Lipman, D.J., Ostell, J. and Sayers, E.W. (2013) GenBank. *Nucleic Acids Res.*, **41**, D36-D42.
32. Madeira, F., Park, Y.M., Lee, J., Buso, N., Gur, T., Madhusoodanan, N., Basutkar, P., Tivey, A.R.N., Potter, S.C., Finn, R.D. *et al.* (2019) The EMBL-EBI search and sequence analysis tools APIs in 2019. *Nucleic Acids Res.*, **47**, W636-W641.
